# miRNA activity contributes to accurate RNA splicing in *C. elegans* intestine and body muscle tissues

**DOI:** 10.1101/479832

**Authors:** Kasuen Kotagama, Anna L Schorr, Hannah S Steber, Marco Mangone

## Abstract

MicroRNAs (miRNAs) are known to modulate gene expression, but their activity at the tissue-specific level remains largely uncharacterized. In order to study their contribution to tissue-specific gene expression, we developed novel tools to profile miRNA targets in the *C. elegans* intestine and body muscle.

We validated many previously described interactions, and identified ~3,500 novel targets. Many of the miRNA targets curated are known to modulate the functions of their respective tissues. Within our datasets we observed a disparity in the use of miRNA-based gene regulation between the intestine and body muscle. The intestine contained significantly more miRNA targets than the body muscle highlighting its transcriptional complexity. We detected an unexpected enrichment of RNA binding proteins targeted by miRNA in both tissues, with a notable abundance of RNA splicing factors.

We developed *in vivo* genetic tools to validate and further study three RNA splicing factors identified as miRNA targets in our study (*asd-2*, *hrp-2* and *smu-2*), and show that these factors indeed contain functional miRNA regulatory elements in their 3’UTRs that are able to repress their expression in the intestine. In addition, the alternative splicing pattern of their respective downstream targets (*unc-60*, *unc-52*, *lin-10* and *ret-1*) is dysregulated when the miRNA pathway is disrupted.

A re-annotation of the transcriptome data in *C. elegans* strains that are deficient in the miRNA pathway from past studies supports and expands on our results. This study highlights an unexpected role for miRNAs in modulating tissue-specific gene isoforms, where post-transcriptional regulation of RNA splicing factors associates with tissue-specific alternative splicing.

## INTRODUCTION

Multicellular organisms have evolved complex forms of gene regulation achieved at different stages throughout development, and equally executed at pre-, co-, and post-transcriptional levels. Alternative splicing, which leads to the production of different protein isoforms using single mRNA precursors, fine tune these regulatory networks, and contributes to the acquisition of tissue identity and function. In humans, more than 95% of genes undergo alternative splicing (Pan *et al.* 2008; Wang *et al.* 2008), and this mechanism is required to ensure that each tissue possesses the correct gene expression pattern needed to thrive (Baralle and Giudice 2017). Many aberrant alternative splicing events are linked to diseases (Scotti and Swanson 2016; Montes *et al.* 2019).

While several tissue-specific splicing factors are known to directly promote RNA splicing, most of the alternative splicing events are achieved through differential expression of particular classes of RNA binding proteins (RBPs), which in turn bind specific *cis*-acting elements located within exon/intron junctions in a combinatorial manner, promoting or inhibiting splicing. Serine Arginine (SR) proteins recognize exon splicing enhancers (ESEs) and are important in promoting constitutive and alternative pre-mRNA splicing, while heterogeneous nuclear ribonucleoproteins (hnRNPs) are a large class of nuclear RBPs that bind exon splicing silencers (ESSs) and usually promote exon retention (Matlin *et al.* 2005). The relative expression levels of members from these two classes of splicing factors vary between tissues, and this imbalance is believed to promote the outcome of tissue-specific alternative splicing events (Caceres *et al.* 1994; Zhu *et al.* 2001).

Tissue identity is also achieved through post-transcriptional gene regulation events, mostly occurring through 3′ Untranslated Regions (3′UTRs), which are portions of genes located between the STOP codon and the poly(A) tail of mature eukaryotic mRNAs. 3′UTRs have been recently subjected to intense study as they were found to be targeted by a variety of factors, which recognize small regulatory elements in these regions and are able to modulate the dosage of gene output at the post-transcriptional level (Matoulkova *et al.* 2012; Oikonomou *et al.* 2014; Mayr 2017). While these regulatory mechanisms are still poorly characterized, and the majority of functional elements remain unknown, disorders in the 3′ end processing of mRNAs have been found to play key roles in the loss of tissue identity and the establishment of major diseases, including neurodegenerative diseases, diabetes, and cancer (Conne *et al.* 2000; Mayr and Bartel 2009; Delay *et al.* 2011; Rehfeld *et al.* 2013).

3′UTRs are frequently targeted by a class of repressive molecules named microRNAs (miRNAs). miRNAs are short non-coding RNAs, ~22nt in length, that are incorporated into a large protein complex named the microRNA-induced silencing complex (miRISC), where they guide the interaction between the miRISC and the target mRNA by base pairing, primarily within the 3′UTR (Bartel 2009). The final outcome miRNA targeting can be context-dependent, however mRNAs targeted by the miRISC are typically held in translational repression prior to degradation of the transcript (Ambros and Ruvkun 2018; Bartel 2018). Initial studies showed that although mismatches between miRNAs and their targets are common, many interactions make use of perfectly complementary at a small conserved heptametrical motif, located at position 2-7 at the 5′end of the miRNA (seed region), (Ambros and Ruvkun 2018; Bartel 2018). Later findings showed that while important, the seed region may also contain one or more mismatches while pairing with its target mRNA, and that this element alone is not a sufficient predictor of miRNA targeting (Ha *et al.* 1996; Reinhart *et al.* 2000; Didiano and Hobert 2006; Grimson *et al.* 2007). Compensatory base pairing at the 3′ end of the miRNA (nucleotides 10-13) can also play a role in target recognition (Shin *et al.* 2010; Chi *et al.* 2012), and have been implicated in conferring target specificity to miRNAs that share the same seed regions (Broughton *et al.* 2016; Wolter *et al.* 2017).

miRNAs and their 3′UTR targets are frequently conserved and play a variety of roles in modulating fundamental biological processes across metazoans. Bioinformatic algorithms, such as miRanda (Betel *et al.* 2008), TargetScan (Lewis *et al.* 2005) and PicTar (Lall *et al.* 2006), use evolutionary conservation and thermodynamic principles to identify miRNA target sites, and are the preferred tools for miRNA target identification. Based on these algorithms it was initially predicted that each miRNA controls hundreds of gene products (Chen and Rajewsky 2007). Recent high-throughput wet bench approaches, have validated and expanded on these initial predictions, and provide further evidence that miRNAs can indeed target hundreds of genes, and regulate molecular pathways throughout development and in diseases (Selbach *et al.* 2008; Helwak *et al.* 2013; Wolter *et al.* 2014; Brown *et al.* 2017; Wolter *et al.* 2017).

In the past few years, several groups produced tissue-specific miRNA localization data in mouse, rat, and human tissues (Eisenberg *et al.* 2007; Landgraf *et al.* 2007) and in cancer (Jima *et al.* 2010). A previous low-throughput study has identified hundreds of *C. elegans* intestine and muscle specific miRNAs and their targets, which are mostly involved in the immune response to pathogens (Kudlow *et al.* 2012). This study used a microarray-based approach, which unfortunately does not provide enough depth to fully understand miRNA function in a tissue-specific manner. In addition, this studies identified only a subset of miRNA targets, which rely on the scaffolding proteins AIN-1 and AIN-2, later found to be only present at specific developmental stages (Kudlow *et al.* 2012; Jannot *et al.* 2016).

In *C. elegans* there are three known Argonaute proteins that execute the miRNA pathway, and are named *alg-1*, *alg-2* and *alg-5*. A recent transcriptome analysis in strains deficient in each of these members show a remarkable difference in function, where *alg-1* and *alg-2* are mostly expressed in somatic tissue and are functionally redundant, and *alg-5* is expressed exclusively in the gonads, interacts with only a subset of miRNAs and is required for optimal fertility (Brown *et al.* 2017). A more recent study used a novel methylation-dependent sequencing approach (mime-Seq), and identified high quality tissue-specific miRNAs in intestine and body muscle tissues (Alberti *et al.* 2018).

Taken together, these studies unequivocally show that there are indeed distinct functional miRNA populations in tissues, which are in turn capable of reshaping transcriptomes and contributing to the acquisition and maintenance of cell identity. Since most miRNAs targets are only predicted, it is still unclear how these events are initiated and maintained.

Our group has pioneered the use of the round worm *C. elegans* to systematically study tissue-specific gene expression (Blazie *et al.* 2015; Blazie *et al.* 2017). In a previous study, we developed a method to isolate and sequence high quality tissue-specific mRNA from worms, and published several integrative analyses of gene expression in most of the *C. elegans* somatic tissues, including the intestine and body muscle (Blazie *et al.* 2015; Blazie *et al.* 2017). In these studies, we found an abundance of several tissue-specific RNA splicing factors, which could explain tissue-specific alternative splicing events. For example, we detected the RNA splicing factors *asd-2* and *sup-12*, previously shown to regulate splicing patterns of the *unc-60* gene in the *C. elegans* body muscle (Ohno *et al.* 2012), and *hrp-2*, a hnRNP known to induce alternative splicing isoforms of the three widely expressed genes; *unc-52* and *lin-10* and *ret-1* (Kabat *et al.* 2009; Heintz *et al.* 2017). The human orthologues of *hrp-2*, hnRNPQ and hnRNPR, have been shown to act in a dosage dependent manner to regulate the alternative splicing of the widely expressed gene PKM, demonstrating the importance of regulating the dosage of hnRNPs (Chen and Cheng 2012). Studies performed using human cell lines have revealed that miRNA-based regulation of splicing factor dosage can drive tissue development (Makeyev *et al.* 2007).

In order to better understand the tissue-specific contribution of miRNA-based regulation to, gene dosage, RBP’s functions, and tissue identity, we performed RNA immunoprecipitation of the *C. elegans* Argonaute ortholog *alg-1*, isolated and sequenced the tissue-specific targets of miRNAs in *C. elegans*, and used them to identify miRNA targets from two of its largest and most well characterized tissues, the intestine and body muscle.

As expected, we found that the number of genes regulated in each tissue correlates with its transcriptome size. However, there is a greater proportion of the transcriptome regulated in the intestine when compared to the body muscle, suggesting that the degree of regulation by miRNA in tissues is heterogeneous. In addition, a large number of identified targets possess RNA binding domains and include several mRNA splicing factors such as hnRNPs and SR proteins. We also detected and validated several tissue-specific miRNA-based regulatory networks involved in tissue-specific alternative splicing of genes. The analysis of splice junctions in transcriptomes from miRNA-deficient strains from past studies support and expand these observations, suggesting a potential role for miRNA in regulating mRNA biogenesis in addition to mRNA turnover.

## MATERIALS and METHODS

### Preparing MosSCI vectors for generating GFP::ALG-1 strains

The strains used for the ALG-1 pull-down were prepared using a modified version of the previously published polyA-pull construct (Blazie *et al.* 2015; Blazie *et al.* 2017). We produced a second-position Entry Gateway vector containing the genomic sequence of *alg-1* tagged at its N-terminus with the GFP fluorochrome. Briefly, we designed primers flanking the coding sequence of *alg-1* and performed a Polymerase Chain Reaction (PCR) amplification to clone the *alg-1* locus from genomic DNA extracted from N2 *wt* worms (primer 1 and 2 in Table S2). The resulting PCR product was analysed on a 1% agarose gel, which displayed a unique expected band at ~3,500 nucleotides. This band was then isolated using the QIAquick Gel Extraction Kit (QIAGEN, cat. 28704) according to the manufacturer’s protocol. Upon recovery, we digested the purified PCR product with the restriction enzymes SacI and BamHI and then cloned it into the modified polyA-pull construct (Blazie *et al.* 2015; Blazie *et al.* 2017), replacing the gene *pab-1*. The ligation reaction was performed using the NEB Quick Ligation Kit (cat. MS2200S) according to the manufacturer’s protocol. We used the QuikChange II Site-Directed Mutagenesis Kit (Agilent, cat. 200523) to remove the unnecessary C-terminal 3xFLAG tag from the polyA-pull vector (primers 3 and 4 in Table S2). We then cloned the previously described endogenous *alg-1* promoter (Vasquez-Rifo *et al.* 2012) by designing primers to add Gateway BP cloning elements, and then performed PCR using N2 *wt* genomic DNA as a template (primers 5 and 6 in Table S2). Using the resulting PCR product, we performed a Gateway BP cloning reaction into the pDONR P4P1R vector (Invitrogen) according to the manufacturers protocol. To assemble the final injection clones, we performed several Gateway LR Clonase II plus reactions (Invitrogen, cat. 12538-013) using the destination vector CFJ150 (Frokjaer-Jensen *et al.* 2012), the tissue-specific or endogenous promoters (*alg-1* for endogenous, *ges-1* for the intestine and *myo-3* for the body muscle), the *gfp* tagged *alg-1* coding sequence, and the *unc-54* 3′UTR as previously published(Blazie *et al.* 2017).

### Microinjections and screening of transgenic C. elegans strains

To prepare single copy integrated transgenic strains we used the *C. elegans* strain Eg6699 [ttTi5605 II; unc-119(ed3) III; oxEx1578](Frokjaer-Jensen *et al.* 2012), which is designed for MosI mediated single copy integration (MosSCI) insertion, using standard injection techniques. These strains were synchronized by bleaching(Porta-DE-LA-Riva *et al.* 2012), then grown at 20°C for 3 days to produce young adult (YA) worms. YA worms were then picked and subjected to microinjection using a plasmid mix containing; pCFJ601 (50ng/μl), pMA122 (10ng/μl), pGH8 (10ng/μl), pCFJ90 (2.5ng/μl), pCFJ104 (5ng/μl) and the transgene (22.5ng/μl)(Frokjaer-Jensen *et al.* 2008). Three injected worms were isolated and individually placed into single small nematode growth media (NGM) plates (USA Scientific, cat 8609-0160) seeded with OP50-1 and were allowed to grow and produce progeny until the worms had exhausted their food supply. The plates were then screened for progenies that exhibited wild type movement and proper GFP expression, and single worms exhibiting both markers were picked and placed onto separate plates to lay eggs overnight. In order to select for single copy integrated worms, an additional screen was performed to select for worms that lost the mCherry fluorochrome expression (extrachromosomal injection markers).

### Genotyping of transgenic C. elegans strains

Single adult worms were isolated and allowed to lay eggs overnight and then genotyped for single copy integration of the transgene by single worm PCR as previously described (Broughton *et al.* 2016) (primers 7-9 in Table S2). Progeny from worms that contained the single copy integrations were propagated and used for this study. A complete list of worm strains produced in this study is shown in Table S3.

### Validating expression of the transgenic construct

To validate the expression of our transgenic construct, and to evaluate our ability to immunoprecipitate GFP tagged ALG-1, we performed an immunoprecipitation (as described below) followed by a western blot. For the western blot we used a primary anti-GFP antibody (Novus, NB600-303) (1:2000) and a fluorescent secondary antibody (LI-COR, 925-32211)(1:5000), followed by imaging using the ODYSSEY CLX system (LI-COR Biosciences, NE) (Figure S1).

### In vivo validation of GFP::ALG-1 functionality by brood size assay

In order to validate the *in vivo* functionality of our transgenic GFP tagged ALG-1, we used a genetic approach. It was previously shown that the knock out *alg-1* strain RF54 [*alg-1(*gk214) X] lead to a decrease in fertility (Bukhari *et al.* 2012). We rescued this decrease in fertility in the *alg-1* knockout strain RF54[*alg-1(*gk214) X] by crossing it into our strain MMA17 (Table S3), which expresses our GFP tagged transgenic ALG-1, driven by the endogenous *alg-1* promoter. The resulting strain MMA20 [*alg-1*(*gk214*)X; *alg-1p*::*gfp*::*alg-1*::*unc-54* II] only expresses our cloned *alg-1* gene tagged with the GFP fluorochrome. We validated the genotype of MMA20 using single worm PCRs as previously described(Broughton *et al.* 2016) (primers 10 and 11 in Table S2 and Figure S2). The brood size assay was used to evaluate the ability of our transgenic GFP tagged ALG-1 construct to rescue the loss in fertility seen in the *alg-1* knockout strain (RF54). The brood size assay was performed by first synchronizing N2 (*wt*), RF54 and MMA20 strains to arrested L1 larvae, through bleaching followed by starvation overnight in M9 solution. We then plated the L1 arrested worms on NGM plates seeded with OP50-1 and allowed the worms to develop to the adult stage for 48 hours after which single worms were isolated onto OP50-1 seeded plates. The adult worms were left to lay eggs overnight (16 hours) after which the adult worms were removed. The eggs were allowed to hatch and develop for 24 hours and the number of larvae in each plate was counted.

### Sample preparation and crosslinking

0.5ml of mixed stage *C. elegans* of each strain was grown on five large 20 cm plates (USA Scientific, cat 8609-0215) and harvested by centrifugation at 400rcf for 3 minutes. The pellets were initially washed in 15ml dH_2_O water and spun down at 400 rcf for 3 minutes and then resuspended in 10ml of M9+0.1%Tween20 and then cross-linked 3 times on ice, with energy setting: 3000 × 100 μJ/cm^2^ (3kJ/m^2^) (Stratalinker 2400, Stratagene)(Moore *et al.* 2014). After the crosslinking, each *C. elegans* strain was recovered by centrifugation at 400 rcf for 3 minutes, and resuspended in two volumes (1ml) of lysis buffer (150mM NaCl, 25mM HEPES(NaOH) pH 7.4, 0.2mM DTT, 10% Glycerol, 25 units/ml of RNasin^®^ Ribonuclease Inhibitor (Promega, cat N2611), 1% Triton X-100 and 1 tablet of protease inhibitor for every 10ml of lysis buffer (Roche cOmplete ULTRA Tablets, Sigma, cat 5892791001). The lysed samples were subjected to sonication using the following settings: amplitude 40%; 5x with 10sec pulses; 50sec rest between pulses (Q55 Sonicator, Qsonica). After the sonication, the cell lysate was cleared of debris by centrifugation at 21,000rcf at 4°C for 15 min and the supernatants were then transferred to new tubes.

### GFP-TRAP bead preparation and immunoprecipitation

25μl of GFP-TRAP beads (Chromotek, gtma-10) (total binding capacity 7.5μg) per immunoprecipitation were resuspended by gently vortexing for 30 seconds, and washed three times with 500μl of cold Dilution/Wash buffer (10 mM Tris/Cl pH 7.5; 150 mM NaCl; 0.5 mM EDTA). The beads were then resuspended in 100μl/per IP of Dilution/Wash buffer. 100μl of resuspended beads were then incubated with 0.5ml of lysate for 1 hour on the rotisserie at 4°C. At the completion of the incubation step, the beads were collected using magnets. The unbound lysate was saved for PAGE analysis. The beads containing the immunoprecipitatred *alg-1* associated to the target mRNAs were then washed three times in 200μl of Dilution/Wash buffer (10 mM Tris/Cl pH 7.5; 150 mM NaCl; 0.5 mM EDTA), and then the RNA/protein complex was eluted using 200μl of Trizol (Invitrogen, cat 15596026) and incubated for 10 minutes at room temperature.

### Trizol/Driectzol RNA purification

The RNA purification was performed using the RNA MiniPrep kit (Zymo Research, cat ZR2070) as per the manufacturers protocol. All centrifugation steps were performed at 21,000g for 30 seconds. We added an equal volume ethanol (95-100%) to each sample in Trizol and mixed thoroughly by vortexing (5 seconds, level 10). The samples were then centrifuged, recovered using a magnet, and the supernatant was transferred into a Zymo-Spin IIC Column in a Collection Tube and centrifuged. The columns were then transferred into a new collection tube and the flow through were discarded. 400 µl of RNA wash buffer was added into each column and centrifuged. In a separate RNase-free tube, we added 5 µl DNase I (6 U/µl) and 75 µl DNA Digestion Buffer, mixed and incubated at room temperature (20-30°C) for 15 minutes. 400 µl of Direct-zol RNA PreWash (Zymo Research, cat ZR2070) was added to each sample and centrifuged twice. The flow-through was discarded in each step. 700 µl of RNA wash buffer was then added to each column and centrifuged for 2 minutes to ensure complete removal of the wash buffer. The columns were then transferred into RNase-free tubes, and the RNAs were eluted with 30 µl of DNase/RNase-Free Water added directly to the column matrix and centrifuging.

### cDNA library preparation and sequencing

Each cDNA library was prepared using a minimum of 500pg of immunoprecipitated RNA from each tissue. The total RNA was reverse transcribed using the IntegenX’s (Pleasanton, CA) automated Apollo 324 robotic preparation system using previously optimized conditions(Blazie *et al.* 2015).

The cDNA synthesis was performed using a SPIA (Single Primer Isothermal Amplification) kit (IntegenX and NuGEN, San Carlos, CA)(Kurn *et al.* 2005). The cDNA was then sheared to approximately 300 bp fragments using the Covaris S220 system (Covaris, Woburn, MA). We used the Agilent 4200 TapeStation instrument (Agilent, Santa Clara, CA) to quantify the abundance of cDNAs and calculate the appropriate amount of cDNA necessary for library construction. Tissue-specific barcodes were then added to each cDNA library, and the finalized samples were pooled and sequenced using the HiSeq platform (Illumina, San Diego, CA) with a 1x75bp HiSeq run.

### Data analysis

We obtained ~15M unique reads per sample (~130M reads total). The software Bowtie 2 (Langmead *et al.* 2009) run using default parameters was used to perform the alignments to the *C. elegans* genome WS250. We used custom Perl scripts and Cufflinks (Trapnell *et al.* 2010) algorithm to study the differential gene expression between our samples. A summary of the results is shown in (Figure S3). Mapped reads were further converted into a bam format and sorted using SAMtools software run with generic parameters (Li *et al.* 2009), and used to calculate Fragments Per Kilobase Million (FPKM) values, as an estimate of the abundance of each gene per sample. We used an FPKM ≥ 1 on the median from each replicate as a threshold for identifying positive hits. This stringent approach discarded ~50-75% of mapped reads for each sample (Figure S3B). The quality of our finalized list of target genes was tested using a principle component analysis versus our N2 *wt* negative control (Supplemental Fig S3C).

### Molecular cloning and assembly of the expression constructs

The promoters of candidate genes were extracted from genomic DNA using genomic PCR and cloned into Gateway-compatible entry vectors (Invitrogen). We designed Gateway-compatible primers (primers 12-19 in Table S2) targeting 2,000 bp upstream of a given transcription start site, or up to the closest gene. Using these DNA primers, we performed PCRs on *C. elegans* genomic DNA, amplified these regions, and analysed the PCR products by gel electrophoresis. Successful DNA amplicons were then recombined into the Gateway entry vector pDONR P4P1R using Gateway BP Clonase reactions (Invitrogen). The reporter construct pAPAreg has been previously described in Blazie et al., 2017 (Blazie *et al.* 2017). The coding region of this construct was prepared by joining the coding sequence of the mCherry fluorochrome to the SL2 splicing element found between the *gpd-2* and *gpd-3* genes, and to the coding sequence of the GFP gene. The entire cassette was then PCR amplified with Gateway-compatible primers and cloned into pDONR P221 by Gateway BP Clonase reactions (Invitrogen).

The 3′UTRs of the genes in this study were cloned by anchoring the Gateway-compatible primers at the translation STOP codon of each gene, to the longest annotated 3′UTR. We have included 50 base pairs downstream of the annotated PAS site to include 3′end processing elements (primers 20-27 in Table S2). The PCR products were analysed using gel electrophoresis analysis and used to perform Gateway BP Clonase reactions (Invitrogen, cat. 11789020) into pDONR P2RP3 as per the manufacturers protocol. The *unc-54* 3′UTR used in this study was previously described in Blazie et al., 2017. The constructs injected were assembled by performing Gateway LR reactions (Invitrogen) with each promoter, reporter, and 3′UTR construct per the manufacturers protocol into the MosSCI compatible destination vector CFJ150. We then microinjected each reporter construct (100ng/μl) with CFJ601 (100ng/μl) into MosSCI compatible *C. elegans* strains using standard microinjection techniques (Evans (ED.)).

### Fluorescent imaging and analysis of nematodes

Confocal images used in Figure 4 were acquired in the Biodesign Imaging Core, Division of the Arizona State University Bioimaging Facility. Transgenic strains were grown at room temperature on NGM plates seeded with OP50-1. The mixed stage worms were washed twice with M9 and resuspended in 1mM of levamisole before imaging using a Nikon C1 Ti-E microscope with 488 nm and 561 nm lasers, 0.75 numerical apature, 90 μM pinhole microscope with a 40x magnification objective lens. We acquired 10 images for each transgenic strain (total 40 images) using the same microscope settings. The fluorescence of GFP and mCherry fluorochromes from the acquired images were individually quantified using the integrated density (ID) function of the ImageJ software (Schneider *et al.* 2012). Fluorescence ratios were then calculated for each worm (n=10, total 40 images) by dividing the ID for GFP by the ID for mCherry. The finalized result for each strain is the averaged fluorescence ratio calculated across all 10 imaged worms. We performed a two tailed student t-test to compare the mean fluorescence ratios for each strain with a p-value cut off <0.05 to establish the presence of post-transcriptional gene regulation.

### Bioinformatic analysis of tissue-specific miRNA targeting biases

The tissue-specific miRNA studies were performed in two steps. First, we utilized custom-made Perl scripts to scan across the longest 3′UTR of each *C. elegans* protein coding gene (WS250) in our datasets, searching for perfect sequence complementarity to the seed regions of all *C. elegans* miRNAs present in the miRBase database (release 21) (Griffiths-Jones 2004; Griffiths-Jones *et al.* 2006; Griffiths-Jones *et al.* 2008; Kozomara and Griffiths-Jones 2011; Kozomara and Griffiths-Jones 2014). This result was then used to calculate the percentage of seed presence in the intestine and body muscle datasets. To calculate the percentage of predicted targets, we extracted both predicted target genes, and their target miRNA name from the miRanda database (Betel *et al.* 2008) and compared the results with our study. A complete list of miRNA predictions for each tissue profiled is shown in Table S1.

### Comparison with other datasets

We extracted the WormBase IDs of genes in the intestine and body muscle transcriptomes previously published by our group (Blazie *et al.* 2017), and most abundant miRNA targets (transcript names) identified by Kudlow et al., 2012 in these tissues (Kudlow *et al.* 2012). We then translated the transcript names from Kudlow et al., 2012 into WormBase IDs using custom Perl scripts, and compared how many genes in each of these groups overlap with our ALG-1 pull-downs. The results are shown in Figure S4. For the analysis shown in Figure S6 we extracted the names of the miRNAs previously identified by Alberti et al., 2018 in the *C. elegans* intestine and body muscle tissues (Alberti *et al.* 2018). We then used custom Perl scripts to search for the presence of the seed regions of these miRNAs in the 3’UTRs of the genes identified in this study (Figure S6).

### Re-annotation of alg-1 and alg-2 knockout transcriptome datasets and splice junction identification

We downloaded from the GEO database the following transcriptome datasets published by Brown et al., 2017 (Brown *et al.* 2017): Project number GSE98935, Wild type Rep 1-3 (GSM2628055, GSM2628056, GSM2628057); *alg-1*(*gk214*) Rep 1-3 (GSM2628061, GSM2628062, GSM2628063); *alg-2*(*ok304*) Rep 1-3 (GSM2628064, GSM2628065, GSM2628066). We used in-house Perl scripts to prepare the reads for mapping, and then these reads as input to the TopHat algorithm (Trapnell *et al.* 2012) to map splice junctions in all nine datasets independently. The TopHat algorithm mapped between 30-56M reads to splice junctions in each sample. *wt*_rep1; 43,721,355 mapped reads (64% of total input reads), *wt*_rep2; 44,440,441 (64%), *wt*_rep3; 37,248,408(62.7%), *alg-1*_rep1; 30,808,645 (62.3%), *alg-1*_rep2; 35,914,514 (63.2%), *alg-1*_rep3; 43,721,355(63.9%), *alg-2*_rep1; 54,471,761(63.2%), *alg-2*_rep2; 56,000,173 (66.8%), *alg-2*_rep3; 46,638,369 (63.9%). We then combined the mapped reads obtained in the three replicates for each strain and used the open source software regtools (Griffith Lab, McDonnell Genome Institute) to annotate these splice junctions using the following command ‘regtools junctions annotate junctions.bed WS250.fa WS250.gtf’. The software produced ~41.8k splice junctions supported by at least 10 reads for the combined N2 *wt* dataset, ~42.3k splice junctions for the *alg-1* dataset and 46.3k for the *alg-2* dataset. We analysed the three resulting cumulative datasets normalized by dividing each score by the total number of mapped reads within each sample. This approach produced 36.7k high quality splice junction for the combine N2 *wt* dataset, ~37k for the *alg-1* combined dataset and ~38k for the combined *alg-2* dataset. The analysis in Figure 6A was performed using splice junctions that are present in all three datasets (30,115 total). To calculate the fold-change for each splice junction, we divided the normalized scores of each splice junction in the *alg-1* and *alg-2* combined datasets by the corresponding scores in the wild type combined dataset. The fold change of each splice junction was then plotted on a log_2_ scale shown in Figure 6.

### RNAi experiments

The RNAi experiments shown in Figure 5 and Figure S7 were performed as follows. N2 worms were synchronized by bleaching and starving overnight in M9 buffer until they reached the L1/dauer stage and then transferred to agar plates containing OP50-1 bacteria, HT115 bacteria with pL4440 *hrp-2* RNAi or pL4440 *asd-2* RNAi (Kamath and Ahringer 2003). We used *par-2* RNAi as a positive control for the experiments, which results in 100% embryonic lethality. To measure the brood size, individual synchronized young adult worms were left overnight (16 hours) to lay eggs. Hatched larvae were counted 24 hours later. Total RNA was extracted from N2 worms treated with either *hrp-2* or *asd-2* RNAi at the adult stage in triplicates.

### RNA extraction for detection of intestine specific splicing variants

We extracted total RNA using the Direct-zol™ RNA MiniPrep Plus kit (Zymo Research, cat ZR2070) from (1) N2 *wt* worms, (2) RF54 (*alg-1(*gk214) X) strain, (3) WM53 (*alg-2*(ok304) II) strain, (4) N2 strain subjected to RNAi as previously described(AHRINGER (ED.)) for *asd-2* and *hrp-2*, and (5) transgenic worms overexpressing the *asd-2* 3′UTR or the *hrp-2* 3′UTR under control of an intestinal promoter (*ges-1p*::pAPAreg::3′UTR). Each strain was synchronized by growing in M9 media to L1/dauer stage then transferred to plates containing HT115. We extracted RNA 48 hours later from adult worms in triplicate for each condition.

### cDNA preparation, image acquisition and splicing isoform analysis

At the completion of the RNA extractions, the cDNA was synthesized from each sample using SuperScript III RT (Life Technologies, cat 18080093) according to the manufacturers protocol. Briefly, 200ng of each RNA sample was incubated with 1 µL of 50mM poly dT anchor, 1 µL of 10mM dNTP mix and brought to a total volume of 14 µL with nuclease free H_2_O and incubated for 5 minutes at 60°C then iced for 1 minute. 4 µL of 5x first strand buffer, 1 µL of 0.1M DTT and 1 µL (200 units) of SuperScript III reverse transcriptase were added to each sample and incubated at 50°C for 60 minutes then heat inactivated at 70°C for 15 minutes. 200ng of cDNA from each sample was used in PCRs consisting of 34 cycles using HiFi Taq Polymerase (Invitrogen, cat 11304011) according to manufacturer protocols. Primers used to test alternative splicing of *unc-60, unc-52, lin-10*, and *ret-1* were designed to flank the alternatively spliced exons and were adapted from previous studies (Kabat *et al.* 2009; Ohno *et al.* 2012; Heintz *et al.* 2017)(Table S2 primers 28-36). We then acquired images of the PCR amplicons (5 µL) separated by agarose gel electrophoresis and assign the alternatively spliced isoforms using the ImageJ software package (Schneider *et al.* 2012). We used the integrated density function of ImageJ by defining equally sized regions of interest around each band in the images and compared the integrated density values by normalizing the smaller bands to the larger bands. The resulting isoform ratios are displayed in Figure S8. Each strain was quantified in triplicate and subjected to a two-tailed student t-test. Statistical significance was assigned for p-values <0.05.

### Data Access

Raw reads were submitted to the NCBI Sequence Read Archive (http://trace.ncbi.nlm.nih.gov/Traces/sra/). The results of our analyses are available in Excel format as Supplementary Table S1, and in our APA-centric website www.APAome.org.

## RESULTS

### A method for the identification of tissue-specific miRNA targets

In order to study the contribution of miRNA activity in producing and maintaining tissue identity, we performed RNA immunoprecipitations of miRNA target genes in two of the largest, morphologically different, and most well characterized tissues in *C. elegans*: the intestine (MCGhee) and body muscle (Gieseler *et al.*) (Figure 1A). We took advantage of the ability of the Argonaute protein to bind miRNA target genes, and cloned *alg-1*, one of the worm orthologs of the human Argonaute 2 protein, downstream of the green fluorescent protein (GFP). The expression of this construct was then driven by the endogenous promoter (*alg-1p*), or restricted to the intestine (*ges-1p*) or body muscle (*myo-3p*) using tissue-specific (TS) promoters (Figure 1B).

**Figure 1:**
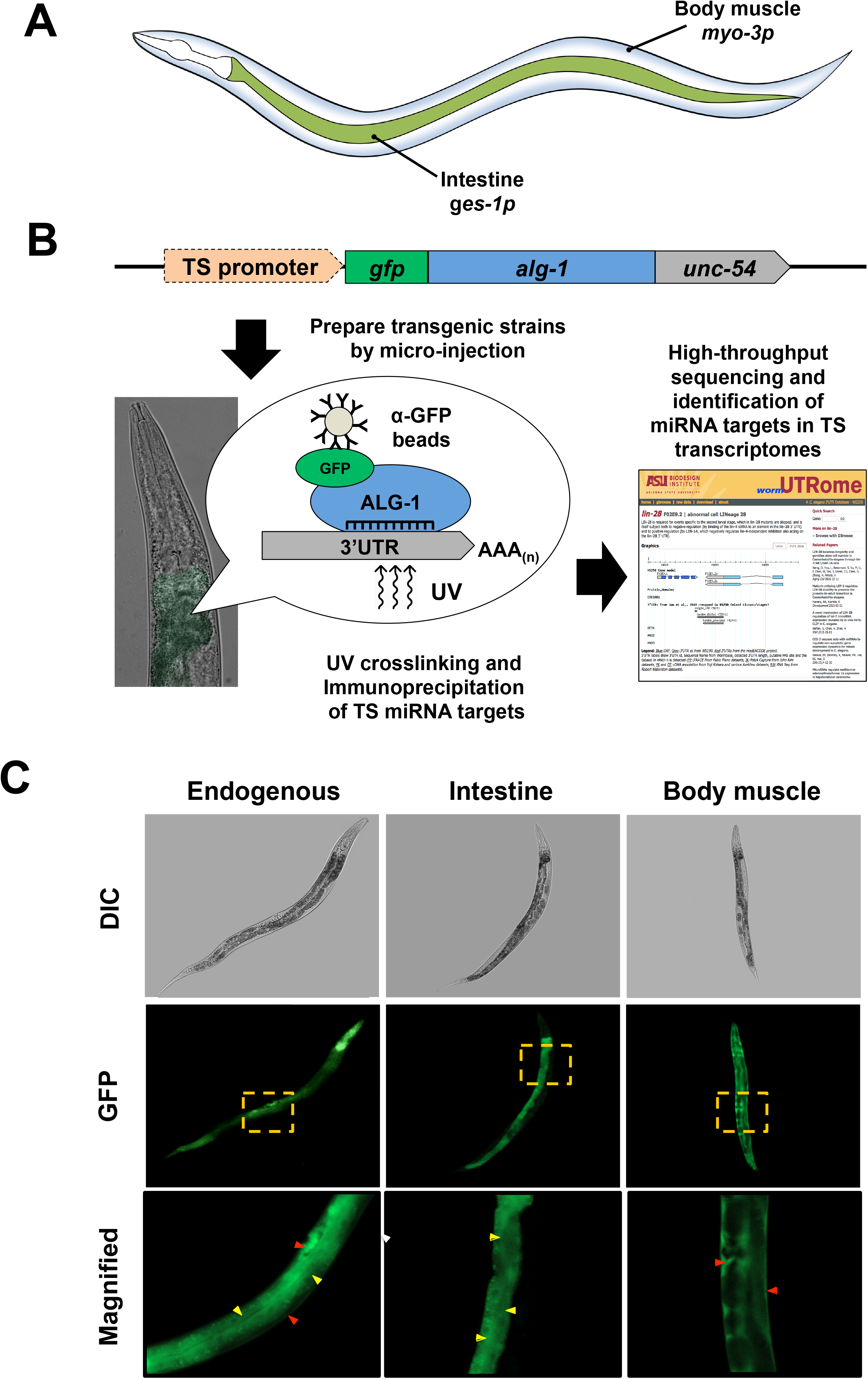
Identification of miRNA targets by tissue-specific immunoprecipitation and sequencing. (A) The anatomical location of the two somatic tissues used in this study. (B) Workflow for the identification of tissue-specific miRNA targets. We cloned the *C. elegans* Argonaute 2 ortholog *alg-1* and fused with the GFP fluorochrome and the unspecific *unc-54* 3′UTR. The expression of this cassette was driven in the intestine and body muscle by using tissue-specific (TS) promoters. These constructs were microinjected into MosSCI-compatible *C. elegans* strains to produce single-copy integrated transgenic animals. These strains were then subjected to UV crosslinking and lysed by sonication. The resulting lysate was subjected to RNA immunoprecipitations with α-GFP antibodies. The resultant tissue-specific miRNA target transcripts were purified, the cDNA libraries were made and sequenced using Illumina HiSeq. (C) Representative images of *C. elegans* single copy integrated strains showing the expression of GFP tagged *alg-1* in endogenous (*alg-1p*), intestine (*ges-1p*), body muscle (*myo-3p*) tissue. Yellow box indicated magnified regions, yellow arrows mark intestine cells, and red arrows mark body muscle cells.

We produced transgenic strains for each construct (Figure 1C) using single copy integration technology (MosSCI) (Frokjaer-Jensen *et al.* 2012; Frokjaer-Jensen *et al.* 2014) to minimize the expression mosaics produced by repetitive extrachromosomal arrays. The strains were validated for integration using genomic PCRs and Western blots (Figure S1).

We then examined the functionality of our cloned *alg-1* in rescue experiments using the *alg-1*^-/-^ strain RF54(*gk214*). This strain has a decrease in fertility caused by the loss of functional *alg-1* (Bukhari *et al.* 2012), which was fully rescued by our cloned *alg-1* construct in a brood size assay (Figure S2), suggesting that our cloned *alg-1* is functional and able to fully mimic endogenous *alg-1*.

We then used our strains to perform tissue-specific RNA immunoprecipitations. Each tissue-specific ALG-1 IP and control IPs were performed in duplicate using biological replicates (total 6 sequencing runs). We obtained ~25M reads on average for each tissue, of which ~80% were successfully mapped to the *C. elegans* genome (WS250) (Figure S3). In order to maximize our success we used very stringent filters to determine gene presence, using only the top 25-50% of genes mapped in each dataset (Figure S3B-C) (Materials and Methods) (Blazie *et al.* 2015; Blazie *et al.* 2017). Our analysis resulted in 3,681 different protein-coding genes specifically targeted by the miRISC using the endogenous *alg-1* promoter or in the intestine or body muscle. The complete list of genes detected in this study is shown in Table S1.

There are only 27 validated *C. elegans* miRNA-target interactions with strong evidence reported in the miRNA target repository miR-TarBase v7, and our study confirmed 16 of these interactions (59%), which is threefold enrichment when compared to a random dataset of similar size (p<0.05, chi square test) (Figure 2A left panel). When compared to genes present in the *C. elegans* intestine and body muscle transcriptomes (Blazie *et al.* 2017), 81% of the intestine and 56% of the body muscle targets identified in this study match with their respective tissues (Figure 2A right panel). A comparison between our hits and a previously published ALG-1 IP dataset in all tissues also support our results (Figure S4) (Zisoulis *et al.* 2010).

**Figure 2:**
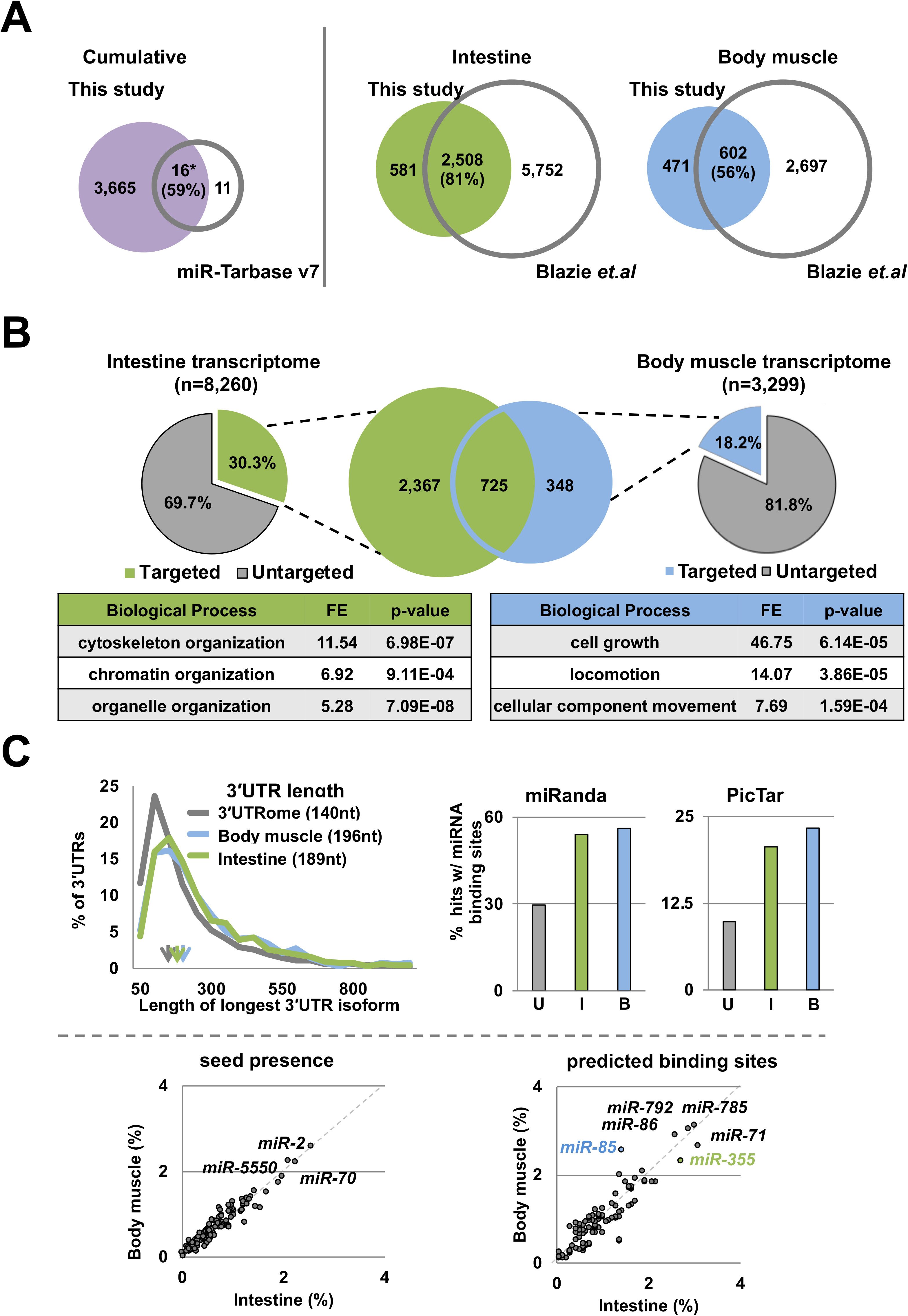
A comparative analysis of ALG-1 immunoprecipitations to identify miRNA targets in the intestine and body muscle. (A) Comparison of miRNA targets identified in this study to other previously published datasets. The numbers indicate protein-coding genes targeted by miRNA in each tissue. Left Panel: Venn diagram showing the comparison of the genes identified in this study to miR-TarBase v7, a compendium of all experimentally validated miRNA targets. 59% of the genes in this database match those identified in this study. Right Panel: Venn diagram showing the comparison of the genes identified in this study to previously published tissue-specific intestine and body muscle transcriptomes. The majority of the targets in both datasets were previously assigned to each tissue. Green – Intestine (*ges-1p*). Blue – Body muscle (*myo-3p*). (B) Pie chart showing the proportion of miRNA targets detected in this study compared to tissue-specific transcriptomes previously characterized by Blazie et.al 2017. The Venn diagram in the center shows the number of protein-coding genes identified is this study as miRNA targets between the intestine and body muscle. The tables show a Gene Ontology analysis for pathway enrichment using the top 100 genes from each dataset used in this study. Green – Intestine (*ges-1p*). Blue – Body muscle (*myo-3p*). (C) Top Left panel: The length of 3′ UTRs from protein coding genes as from the 3′UTRome v1 (Mangone *et al.* 2010) compared to the intestine and body muscle targets identified in this study. The arrows indicate the median 3′UTR length. Genes targeted in the intestine and body muscle have longer 3′UTRs on average than those published in the *C. elegans* 3′UTRome v1. Top Right panel: Proportion of 3′UTRs with predicted miRNA binding sites (miRanda and PicTar) (Lall *et al.* 2006; Betel *et al.* 2008) in the 3′UTRome (gray), and in 3’UTRs of genes identified in this study. Green – Intestine; Blue – Body muscle. The genes identified in this study in the intestine and body muscle are enriched for predicted miRNA binding sites. Bottom panel: Analysis of miRNA target sites identified in this study. The two axis show the proportion of 3′UTRs with perfect seeds or with predicted target sites (miRanda) (Betel *et al.* 2008), normalized to the total number of genes targeted in each tissue for each miRNA. miRNAs that target more than 2% of the genes are listed. The blue mark denotes *miR-85*, a body muscle specific miRNA. The green mark denotes *miR-355*, an intestine specific miRNA.

In order to further validate the quality of our hits, we used GFP-based approaches to confirm the tissue localization of a few tissue-specific genes identified in our study, and found with the exception of one, their observed localization match the expected tissue (Figure S5). In addition, to further test the quality of our data, we compared our results with the intestine and body muscle specific miRNA localization data from past studies (Alberti *et al.* 2018) (Figure S6). We found that more than 84% of the genes identified in our study possess predicted binding sites in their 3’UTRs for miRNAs detected in each tissue, suggesting strong correlation between our results and Alberti et al., 2018 (Alberti *et al.* 2018) (Figure S6).

### ALG-1 targets in the intestine regulate key metabolic enzymes

The *C. elegans* intestine is composed of 20 cells that begin differentiation early in embryogenesis and derive from a single blastomere at the 8-cell stage (MCGhee). As the primary role of the intestine is to facilitate the digestion and the absorption of nutrients, many highly expressed genes in this tissue are digestive enzymes, ion transport channels and regulators of vesicle transport (MCGhee).

In our intestinal ALG-1 pull-down we identified 3,089 protein-coding genes targeted by miRNAs. 2,367 of these genes were uniquely targeted by miRNAs in this tissue (Figure 2B). As expected, and consistent with the function of the intestine, we find a number of enzymes involved with glucose metabolism, such as *enol-1* an enolase, *ipgm-1* a phosphoglycerate mutase, and 3 out of 4 glyceraldehyde-3-phosphate dehydrogenases (*gpd-1*, *gpd-2* and *gpd-4*). The human orthologue of the *C. elegans* gene *enol-1*, *eno1* has been previously identified as a target of *miR-22* in the context of human gastric cancer (Qian *et al.* 2017). In addition, some of our top hits are the fatty acid desaturase enzymes *fat-1*, *fat-2*, *fat-4* and *fat-6*, which are all involved with fatty acid metabolism, suggesting that these metabolic pathways are subjected to a high degree of regulation in the intestine. All of these genes contain seed elements in their 3′ UTRs (Table S1). Additionally, we find 5 out of 6 vitellogenin genes (*vit-1*, *vit-2*, *vit-3*, *vit-5* and *vit-6*) strongly targeted by miRNAs, with *vit-2* and *vit-6* being the most abundant transcripts in our immunoprecipitation (Table S1). *vit-2* was shown to be targeted by ALG-1 in a previous study (Kudlow *et al.* 2012), and both possess MiRanda (Betel *et al.* 2008; Betel *et al.* 2010) and/or PicTar (Lall *et al.* 2006) predicted binding sites (Table S1). These vitellogenin genes produce yolk proteins and are energy carrier molecules synthesized in the intestine. These yolk proteins are then transported to the gonads and into the oocytes to act as an energy source for the developing embryos (Depina *et al.* 2011). Accordingly, we also find a number of RAB family proteins that are responsible for intracellular vesicular transport (*rab-1*, *rab-6.1*, *rab-7*, *rab-8*, *rab-21*, *rab-35* and *rab-39*).

Several transcription factors were also identified as a miRNA targets in the intestine. *skn-1* is a bZip transcription factor that is initially required for the specification of cell identity in early embryogenesis, and then later plays a role in modulating insulin response in the intestine of adult worms (Blackwell *et al.* 2015). This gene has already been found to be targeted by miRNA in many past studies (Zisoulis *et al.* 2010; Kudlow *et al.* 2012) and contains many predicted miRNA binding sites and seed regions from both MiRanda (Betel *et al.* 2008; Betel *et al.* 2010) and PicTar (Lall *et al.* 2006) prediction software (Table S1). A second transcription factor *pha-4* is expressed in the intestine, where it has an effect on dietary restriction mediated longevity (Smith-Vikos *et al.* 2014). *pha-4* is a validated target of *let-7* in the intestine(Grosshans *et al.* 2005), and along with *skn-1*, is also targeted by *miR-228* (Smith-Vikos *et al.* 2014). Additionally, *pha-4* is targeted by *miR-71* (Smith-Vikos *et al.* 2014).

We also find as a target of miRNA, *die-1* a gene which associated with the attachment of the intestine to the pharynx and the rectum (Heid *et al.* 2001), and the chromatin remodeling factor *lss-4* (let seven suppressor), which is able to prevent the lethal phenotype induced by knocking out the miRNA *let-7* (Grosshans *et al.* 2005). These two genes were also validated by others as miRNA targets (Grosshans *et al.* 2005).

The intestine plays an important role in producing an innate immune response to pathogens. The genes *atf-7*, *pmk-1* and *sek-1* were all identified as targets of miRNAs in this tissue. These three genes act together to produce a transcriptional innate immune response where the transcription factor *atf-7* is activated through phosphorylation by kinases *pmk-1* and *sek-1*. Consistent with our findings, the role of miRNAs in regulating the innate immune response through the intestine and these genes has been reported in multiple studies (Ding *et al.* 2008; Kudlow *et al.* 2012; Sun *et al.* 2016).

### Muscle ALG-1 targets modulate locomotion and cellular architecture

*C. elegans* possess 95 striated body wall muscle cells, which are essential for locomotion (Gieseler *et al.*). Its sarcomeres are composed of thick filaments containing myosin associated with an M-line, and thin filaments containing actin associated with the dense body. The pulling of actin filaments by myosin heads generates force that produces locomotion (Moerman and Williams).

Our ALG-1 pull-down identified 1,047 protein-coding genes targeted by miRNAs in the body muscle tissue (Table S1). Within this group, 348 genes were not present in our intestine dataset, and are specifically restricted to the body muscle tissue (Table S1). Our top hits include genes involved in locomotion, and general DNA maintenance (*grd-5*, *gcc-1*, *gop-2*, etc.) and several with unknown function. Consistent with muscle functions, we detected *mup-2*, which encodes the muscle contractile protein troponin T, *myo-3*, which encodes an isoform of the myosin heavy chain, *dlc-1*, which encodes dynein light chain 1 and F22B5.10, a poorly characterized gene involved in striated muscle myosin thick filament assembly. *mup-2*, *myo-3* and *dlc-1* were all found to be targeted by ALG-1 in previous studies (Zisoulis *et al.* 2010; Kudlow *et al.* 2012). Consistent with muscle function, a GO term analysis of this dataset highlights an enrichment of genes involved in locomotion (Table S1), suggesting a potential role for miRNAs in this biological process.

We also identified numerous actin gene isoforms (*act-1*, *act-2*, *act-3* and *act-4*), which are required for maintenance of cellular architecture within the body wall muscle, and the Rho GTPase *rho-1*, which is required for regulation of actin filament-based processes including embryonic polarity, cell migration, cell shape changes, and muscle contraction (Table S1). Small GTPase are a gene class heavily targeted by miRNAs (Enright *et al.* 2003; Liu *et al.* 2012). The human ortholog of *rho-1* is a known target for *miR-31*, *miR-133*, *miR-155* and *miR-185* (Liu *et al.* 2012).

Importantly, we also found several muscle-specific transcription factors including *mxl-3*, a basic helix-loop-helix transcription factor and K08D12.3, an ortholog of the human gene *ZNF9*. These genes are known to regulate proper muscle formation and cell growth. *mxl-3* is targeted by *miR-34* in the context of stress response (Chen *et al.* 2015). Both genes have been detected in past ALG-1 immunoprecipitation studies (Zisoulis *et al.* 2010).

Our top hit in this tissue is the zinc finger CCCH-type antiviral gene *pos-1*, a maternally inherited gene necessary for proper fate specification of germ cells, intestine, pharynx, and hypodermis(Farley *et al.* 2008). *pos-1* contains several predicted miRNA binding sites in its 3′UTR (Table S1), and based on our GFP reporter validation study is strongly expressed in the body muscle (Figure S4). We also find the KH domain containing protein *gld-1*, the homolog of the human gene *QKI*, which is targeted by *miR-214* (Wu *et al.* 2017), *miR-200c* and *miR-375* (Pillman *et al.* 2018).

### miRNA targeting is more extensive in the intestine than it is in the body muscle

By comparing the percentage of tissue-specific miRNA targets identified in our study to the previously published intestine and body muscle transcriptomes (Blazie *et al.* 2015; Blazie *et al.* 2017), we found that the hits in the intestine are almost twice the number of hits we obtained in the body muscle tissue (30.3% vs 18.2%) (Figure 2B). The length of the 3′UTRs of genes identified as miRNA targets in the intestine and the body muscle tissues are similar when comparing the two tissues, but are on average longer and have more predicted miRNA binding sites than the overall *C. elegans* transcriptome (Figure 2C, top panel). Our results indicate that despite similarity in average 3′UTR length in tissues, the extent of miRNA-based regulatory networks are not similar across tissues. In this specific case, we found that the intestine utilizes miRNA-based gene regulation to a greater extent, when compared with the body muscle.

### MiRNA targets in the intestine and body muscle are enriched for *miR-355*and *miR-85* binding sites

A bioinformatic analysis of the longest 3′UTR isoforms of the targeted genes showed there was no specific requirement for the seed regions in either tissue (Figure 2C bottom left panel). However, the use of predictive software showed that in addition to others, there is an intestine-specific bias for *miR-355* targets (Figure 2C, right panel, green mark). This miRNA is involved in the insulin signaling and innate immunity (Zhi *et al.* 2017), which in *C. elegans* are both mediated through the intestine.

In contrast, we observed an enrichment of targets for the poorly characterized *miR-85* in the body muscle dataset (Figure 2C, right panel, blue mark). These two miRNAs are uniquely expressed in the respective tissues (Martinez *et al.* 2008).

While *miR-85* and *miR-355* were the most abundant and tissue-restricted miRNAs identified in this study, several other miRNAs, including *miR-71, miR-86, miR-785* and *miR-792* were also found highly expressed but less spatially restricted.

### Intestine and body muscle miRNAs target RNA binding proteins

Upon further analysis, we observed an unexpected enrichment of genes containing RNA binding domains in both datasets (Supplemental Table 1). RNA binding proteins are known to play an important role in producing tissue-specific gene regulation by controlling gene expression at both the co-and post-transcriptional levels (Tamburino *et al.* 2013), and out of the ~887 RNA binding proteins (RBPs) defined in *C. elegans* (Tamburino *et al.* 2013), we identified almost half as targets of miRNAs across both tissues (45%).

**Table 1:** Summary of expression pattern, miRNA targets and predicted miRNA binding sites for *asd-2*, *hrp-2* and *smu-2*.

| Gene | Transcriptome |  | miRNAome |  | miRNA predictions |  | Known targets |
| --- | --- | --- | --- | --- | --- | --- | --- |
|  | Body muscle | Intestine | Body muscle | Intestine | miRanda | TargetScan |  |
| ***asd-2*** | Y | Y | — | Y | *miR-86, miR-255, miR-259, miR-785* | *miR-1/796, miR-46/47* | *let-2, unc-60* |
| ***hrp-2*** | Y | Y | — | Y | *miR-58, miR-62, miR-80, miR-81, miR-82, miR-83, miR-84, miR-85, miR-86, miR-90, miR-232, miR-244, miR-259, miR-357, miR-785* | *miR-1018, miR-1821, miR-4809* | *ret-1, unc-52, lin-10* |
| ***smu-2*** | — | Y | — | Y | — | *miR-60-3p, miR-234, miR-230, miR-392, miR-789, miR-792, miR-1020, miR-1818, miR-1828* | *unc-52, unc-73* |

We found that out of the 599 known RBPs present in the intestine transcriptome (Blazie *et al.* 2015; Blazie *et al.* 2017), 380 (64%) were present in our intestine dataset as targets of miRNAs (Figure 3A). This is a notable enrichment when compared to transcription factors and non-RBP genes found in these tissues by Blazie et al. 2017 (Blazie *et al.* 2017), of which only a fraction were identified in our study as miRNA targets (Figure 3A left panel). A similar trend is also present in the body muscle, with 170 (54%) of RBPs identified as miRNA targets (Figure 3A left panel). Importantly, the largest pool of targeted RBPs in both tissues was composed of general factors (GF), such as translation factors, tRNA interacting proteins, ribosomal proteins, and ribonucleases (Figure 3A right panel), suggesting extensive miRNA regulatory networks are in place in this tissue.

**Figure 3:**
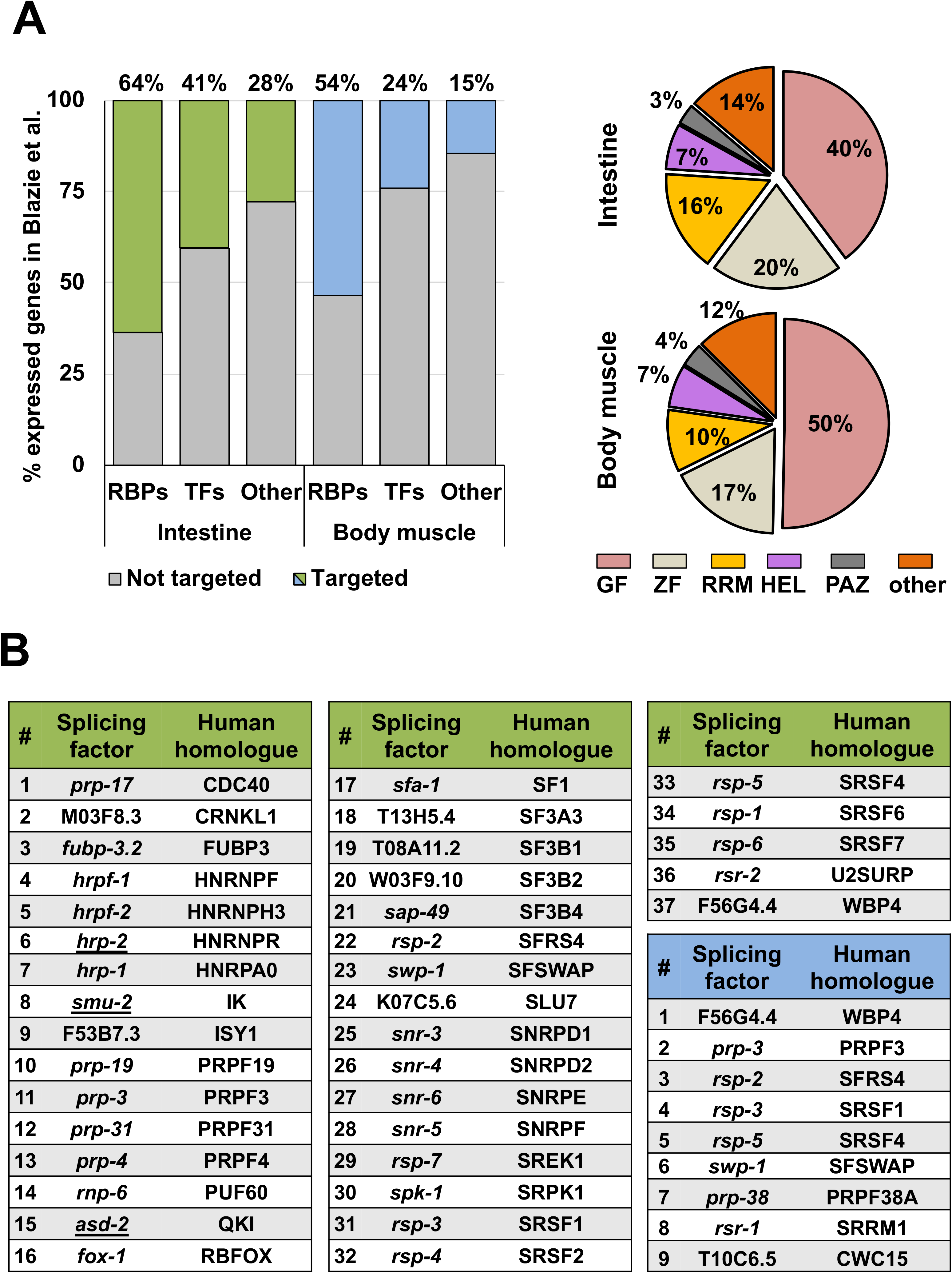
An enrichment of RBPs, and RNA splicing factors targeted by miRNAs in the intestine and body muscle. (A) Left panel: Proportion of RBPs targeted by miRNAs in each tissue. There is an enrichment of RBPs targeted in the intestine (green 63.6%) and the body muscle (blue 53.5%). ‘TFs’ represents genes annotated as transcription factors while ‘Other’ represents protein-coding genes that are not RBPs. Right panel: Subtypes of RBPs targeted by miRNAs in the intestine and body muscle. GF - General Factors, including translation factors, tRNA proteins, ribosomal proteins and ribonucleases; ZF - Zinc finger; RRM - RNA recognition motif; HEL - RNA Helicase; PAZ - PIWI PAZ, PIWI, Argonautes. The majority of the targeted RBPs are general and zinc finger-containing factors. (B) RNA Splicing factors identified as miRNA targets in the intestine (green) and body muscle (blue) tissues. More than half of the RNA splicing factors examined are targeted by miRNAs in the intestine as compared to body muscle.

### miRNAs target RNA splicing factors

A further analysis of the RNA binding proteins targeted in each tissue revealed that one of the most abundant class of RBPs detected in our ALG-1 pull-down in intestine and body muscle datasets was RNA splicing factors (Figure 3B). The *C. elegans* transcriptome contains at least 78 known RNA splicing factors involved in both constitutive and alternative splicing (Tamburino *et al.* 2013). 64 RNA splicing factors (82%) have been previously assigned by our group in the intestine (Blazie *et al.* 2015; Blazie *et al.* 2017) and presumably are responsible for tissue-specific RNA splicing. 31 RNA splicing factors (40%) were also previously assigned by our group to the body muscle tissue (Blazie *et al.* 2015; Blazie *et al.* 2017).

Our tissue-specific ALG-1 pull-down identified 37 RNA splicing factors as miRNA targets in the intestine (~47%) (Figure 3B), and 34 of these were also previously identified by our group as being expressed in this tissue (Blazie *et al.* 2015; Blazie *et al.* 2017). In contrast, we have detected only nine RNA splicing factors targeted by miRNAs in our body muscle tissue ALG-1 pull-down, five of which previously assigned by our group in the body muscle transcriptome (Blazie *et al.* 2015; Blazie *et al.* 2017) (Figure 3B).

The difference in RNA splicing factors targeted by miRNA in these two tissues is significant as with the intestine contains three orders of magnitude more miRNA targeted RNA splicing factors than the body muscle. Of note, many different sub-types of RNA splicing factors identified in this study have human homologs, such as well-known snRNPs, hnRNPs and SR proteins (Figure 3B).

### Expression of the RNA splicing factors *asd-2*, *hrp-2* and *smu-2* is modulated through their 3′UTRs

In order to validate that RNA splicing factors found in our ALG-1 pull-down IPs are targeted by miRNAs in the intestine, we used the pAPAreg dual fluorochrome vector we developed in a past study (Blazie *et al.* 2017) (Figure 4A). This vector uses a single promoter to drive the transcription of a polycistronic pre-mRNA where the coding sequence of the mCherry fluorochrome is separated from the coding sequence of GFP by a SL2 trans-splicing element (SE) (Blazie *et al.* 2017). The test 3′UTR is cloned downstream of the GFP gene. Since the mCherry transcript is trans-spliced, it reports transcription activation. The GFP gene instead reports translational activity; since its expression is dictated by the downstream tested 3′UTR. If a given miRNA targets the test 3′UTR, the GFP intensity decreases when compared with an untargeted 3′UTRs (*ges-1*). By comparing the ratio of the mCherry (indicating transcription) to the GFP (indicating translation) fluorochromes, we are able to define the occurrence of post-transcriptional silencing triggered by the tested 3′ UTR (Figure 4A) (Blazie *et al.* 2017).

**Figure 4:**
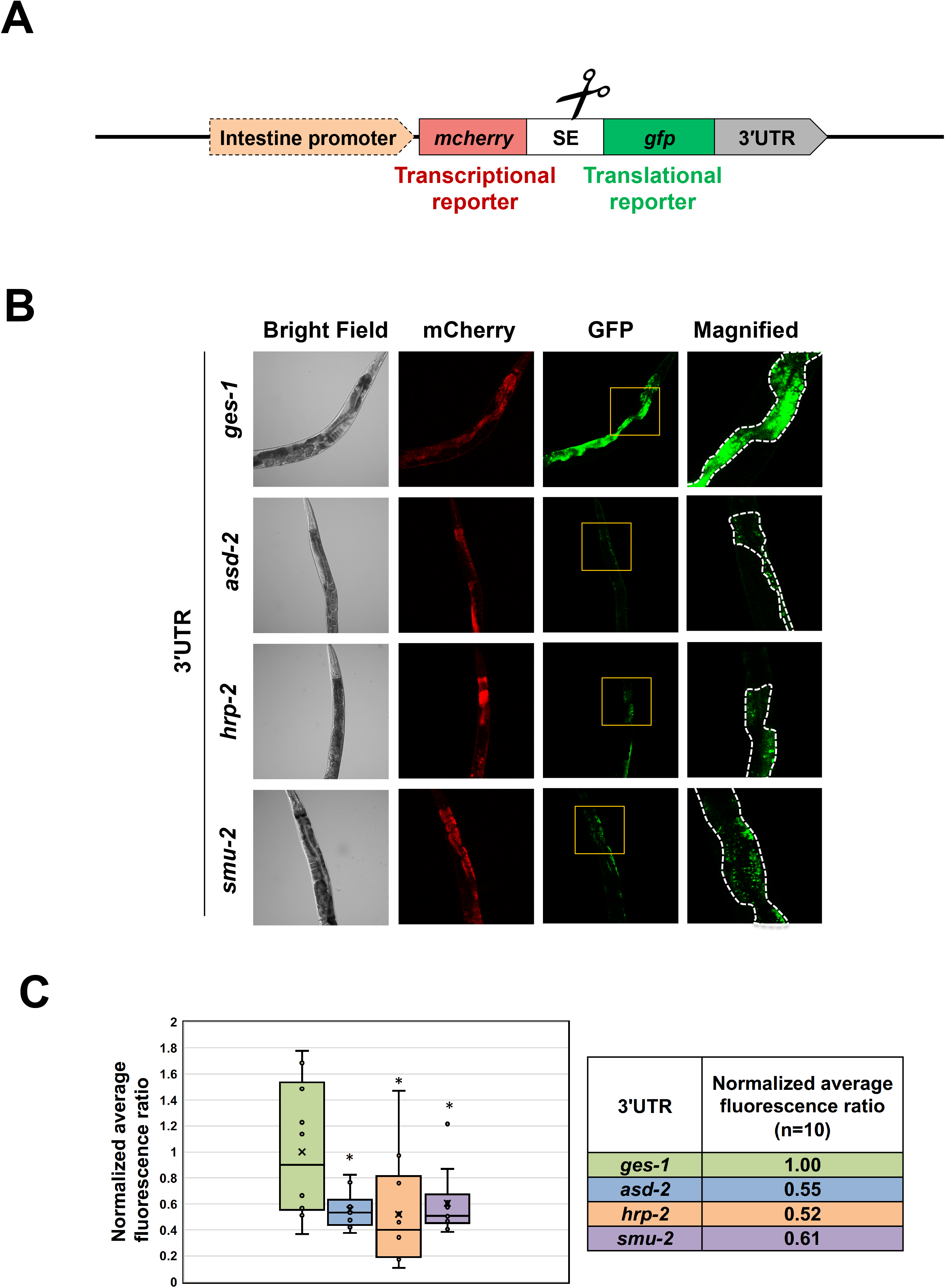
*asd-2*, *hrp-2* and *smu-2* 3′UTRs regulate post-transcriptional gene expression in the intestine. (A) Diagram of the construct used in these experiments (pAPAreg). An intestine-specific promoter drives the expression of a bi-cistronic dual fluorochrome vector in the intestine. The mCherry fluorochrome reports transcription activity of the construct, while the GFP reports post-transcriptional activity through the test 3′UTR cloned downstream of the GFP reporter sequence. If the test 3′UTR is targeted by repressive regulatory factors, such as miRNAs, the GFP fluorochrome lowers in its expression. SE: trans-splicing element extracted from the intergenic region located between the genes *gpd-2* and *gpd-3*. (B) Representative images of *C. elegans* strains generated with pAPAreg constructs expressing one of the following 3′UTRs: *ges-1*, *asd-2*, *hrp-2* or *smu-2* downstream of the GFP fluorochrome. Yellow boxes indicate magnified regions. White dotted lines indicate the intestine. C) The bar graphs show the quantified and normalized mean fluorescence ratio between the GFP and the mCherry fluorochromes. The mean fluorescence ratio is calculated from 10 worms per strain. The error bars indicate the standard error of the mean. *p<0.05. We observed ~40% reduction in normalized GFP intensity modulated by *asd-2, hrp-2*, and *smu-2* 3′UTRs.

We selected three representative RNA splicing factors identified in our study in the intestine (*asd-2*, *hrp-2 and smu-*2) (Table 1) and prepared transgenic strains to validate their expression and regulation (Figure 4B). We used the *ges-1* 3′UTR as a negative control for miRNA targeting, as it is strongly transcribed and translated in the intestine, with no predicted miRNA binding sites (PicTar), and poorly conserved seed regions (TargetScan), suggesting minimal post-transcriptional gene regulation (Egan *et al.* 1995; Marshall and Mcghee 2001). *ges-1* was not significantly abundant in our intestine ALG-1 pull-down (Table S1). The presence of the *ges-1* 3′UTR in the pAPAreg vector led to the expression of both mCherry and GFP fluorochromes, indicating robust transcription and translation of the construct as expected (Figure 4B).

We then cloned *asd-2*, *hrp-2* and *smu-2* 3′UTRs downstream of the GFP fluorochrome in our pAPAreg vector, prepared transgenic worms expressing these constructs, and studied the fluctuation of the expression level of the GFP fluorochrome in these transgenic strains. All three 3′UTRs were able to significantly lower GFP expression when compared to the control strain with the *ges-1* 3′UTR, with ~40% repression, while the mCherry signal was similar in all strains (Figure 4B). These results suggest that these three RNA binding proteins contain regulatory binding sites within their 3′UTRs potentially able to repress their expression.

### MiRNAs target intestine RNA splicing factors promoting tissue-specific alternative splicing

We then tested changes to tissue-specific alternative splicing in the intestine caused by the STAR protein family member *asd-2*, which regulates the alternative splicing pattern of the gene *unc-60*. *unc-60* is expressed as two alternatively spliced isoforms in a tissue-specific manner (Ohno *et al.* 2012) (Figure 5A); *unc-60a* is expressed predominantly in the body muscle while *unc-60b* is expressed in many other tissues including the intestine (Ohno *et al.* 2012).

**Figure 5:**
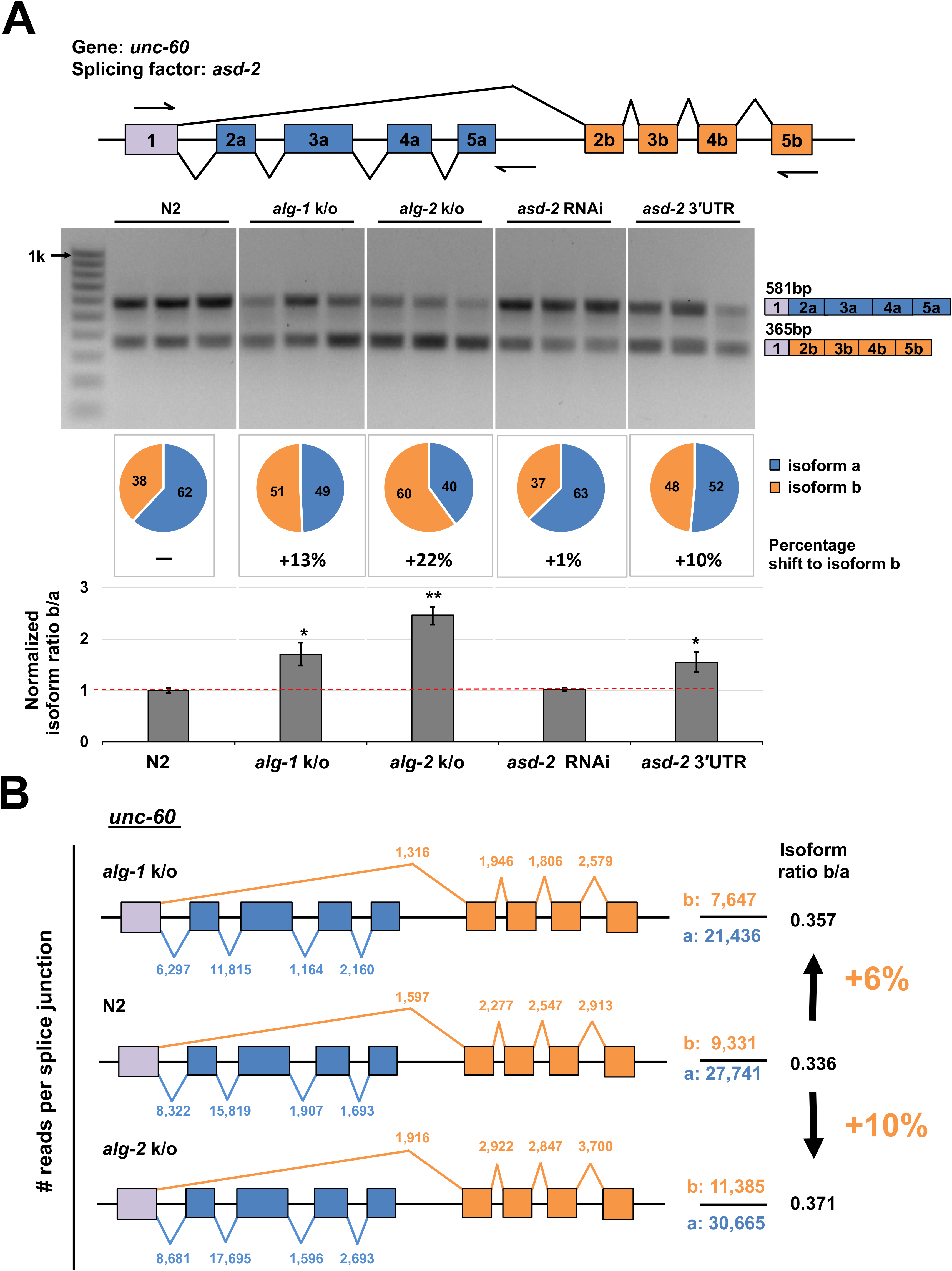
The splicing pattern of *unc-60* is modulated by miRNA activity in the intestine. (A) Top Panel: Schematic of the genomic locus of *unc-60.* This gene is expressed as a longer *unc-60a* isoform, and a shorter *unc-60b* isoform. Arrows mark the binding sites of the primers used to detect the two isoforms. Middle Panel: RT-PCR performed from total RNA extracted from biological replicates in triplicate and visualized in 1% agarose gel. 1) N2: *wt* worms. 2) *alg-1* k/o: RF54[*alg-1*(gk214) X], 3) *alg-2* k/o: WM53[*alg-2*(ok304) II], 4) *asd-2* RNAi: N2 worms subjected to *asd-2* RNAi, 5) Over expression of *asd-2* 3′UTR in the intestine. The pie charts below each gel shows a quantification of each of the occurrence of the two isoforms. The percentage below the pie chart is the increase in *unc-60* isoform b abundance when compared to N2(*wt*). The bar chart shows the change in isoform ratio between strains. The y-axis shows the abundance ratio (shorter isoform/longer isoform) of the two alternatively spliced isoforms examined. Exon skipping increases in *alg-1* and *alg-2* k/o strains, and in *asd-2* 3′UTR overexpression strains. Error bars indicate standard error of the mean. Student t-test *p<0.05 **p<0.01. (B) A comparison of the splice junction usage in *unc-60* as observed in transcriptome data for *alg-1* and *alg-2* knockout strains (Brown *et al.* 2017). The numbers above each slice junction indicates the number of reads mapped to that splice junction. The total reads for each isoform are indicated next to the gene model. The isoform ratios indicated next to the gene models are calculated by dividing the total reads for each isoform. There is a ~6-10% increase in the expression of the shorter *unc-60b* isoform in the miRNA deficient strains. Blue: reads corresponding to *unc-60a.* Orange: reads corresponding to *unc-60b*

We first tested the *unc-60* RNA isoform ratio in *wt* N2 worms. We extracted total RNA from N2 worms in triplicate and performed RT-PCR experiments using primers flanking the two *unc-60* isoforms (Figure 5A). As expected, we found that the *unc-60a* longer isoform was more abundantly expressed in *wt* worms (62%) (Figure 5A).

We then investigated if the miRNA pathway has a role in regulating these splicing events, by testing changes in *unc-60* isoform abundance in the *alg-1* and *alg-2* knockout strains (RF54 (*alg-1*(*gk214*) X) and WM53(*alg-2*(*ok304*) II). These strains are deficient in miRNA-based gene regulation. We found that loss of these miRNA effectors lead to a 10-20% shift in the expression of the two *unc-60* isoforms (Figure 5A), indicating the importance of the miRNA pathway in regulating alternative splicing of this gene.

We then used a genetic approach to test the alternative splicing of this gene in the context of miRNA regulation. We reasoned that if ALG-1 targets the *asd-2* 3′UTR in the intestine lowering the expression of *asd-2*, which in turn causes *unc-60* alternative splicing pattern, we should be able to interfere with this mechanism by overexpressing the *asd-2* 3′UTR in this tissue and in turn test the role of the miRNA pathway in this process.

As expected, the overexpression of the *asd-2* 3′UTR in the intestine led to changes in the *unc-60* alternative splicing pattern, indicating that post-transcriptional regulation of *asd-2* through its 3′UTR is important for the alternative splicing pattern of *unc-60* in the intestine (Figure 5A). Conversely, *asd-2* RNAi did not induce changes in *unc-60* alternative splicing pattern (Figure 5A). We validated the efficiency of our RNAi experiments by performing a brood size assay, which indicated strong RNAi activity (Figure S7). Similar results were observed by testing a second splicing factors (*hrp-2*) known to direct alternative splicing of the genes *ret-1*, *lin-10* and *unc-52* (Kabat *et al.* 2009; Heintz *et al.* 2017) (Figure S8).

### Loss of miRNA function lead to dispersed changes in splice junction usage

Since our data support a role for the miRNA pathway in modulating mRNA biogenesis, we were interested in testing the extent of these effects at thetranscriptome level. We decided to downloaded and mapped splicing junctions in genes from *alg-1* and *alg-2* knockout strains previously published by Brown et al., 2017 (Brown *et al.* 2017). These worm strains are viable but are severely impaired. We reasoned that if the miRNA pathway contributes at some level to mRNA biogenesis, we should be able to see widespread changes in the usage of splice junctions in these datasets. To test this hypothesis, we downloaded the *alg-1* and *alg-2* datasets (three replicates for each strain plus *wt* N2 control), and extracted splice junction information. We first tested if the effects we observed in *unc-60* with our biochemical and genetic approaches (Figure 5A) could be also detected in these datasets. In the case of *unc-60*, there is a 6-10% change in splice junction usage between isoforms consistently in all re-annotated replicates, in both *alg-1* and *alg-2* knockout strains (Figure 5B). This result is in line with our analysis in Figure 5A. A similar and more striking aberrant splice junction usage is observed in the case of *lin-10*, and *unc-52*, and with a less pronounced effect in *ret-1* (Figure S9). These results are also in agreement with our study in Figure S8.

We then expanded this analysis to all splicing junctions we were able to map using these transcriptomes. From a total of 30,115 high quality known splice junctions present in all three datasets (Table S4), we identified ~3,946 of them in ~2,915 protein coding genes that were affected by more than 2-fold change in usage in both *alg-1* and *alg-2* knockout datasets (~13.2% of the total mapped splice junctions) (Figure 6A). In addition, we detect several cases of exon inclusion, skipping and aberrant splicing events that occur exclusively in the *alg-1* and/or *alg-2* mutant strains (Figure 6B).

**Figure 6:**
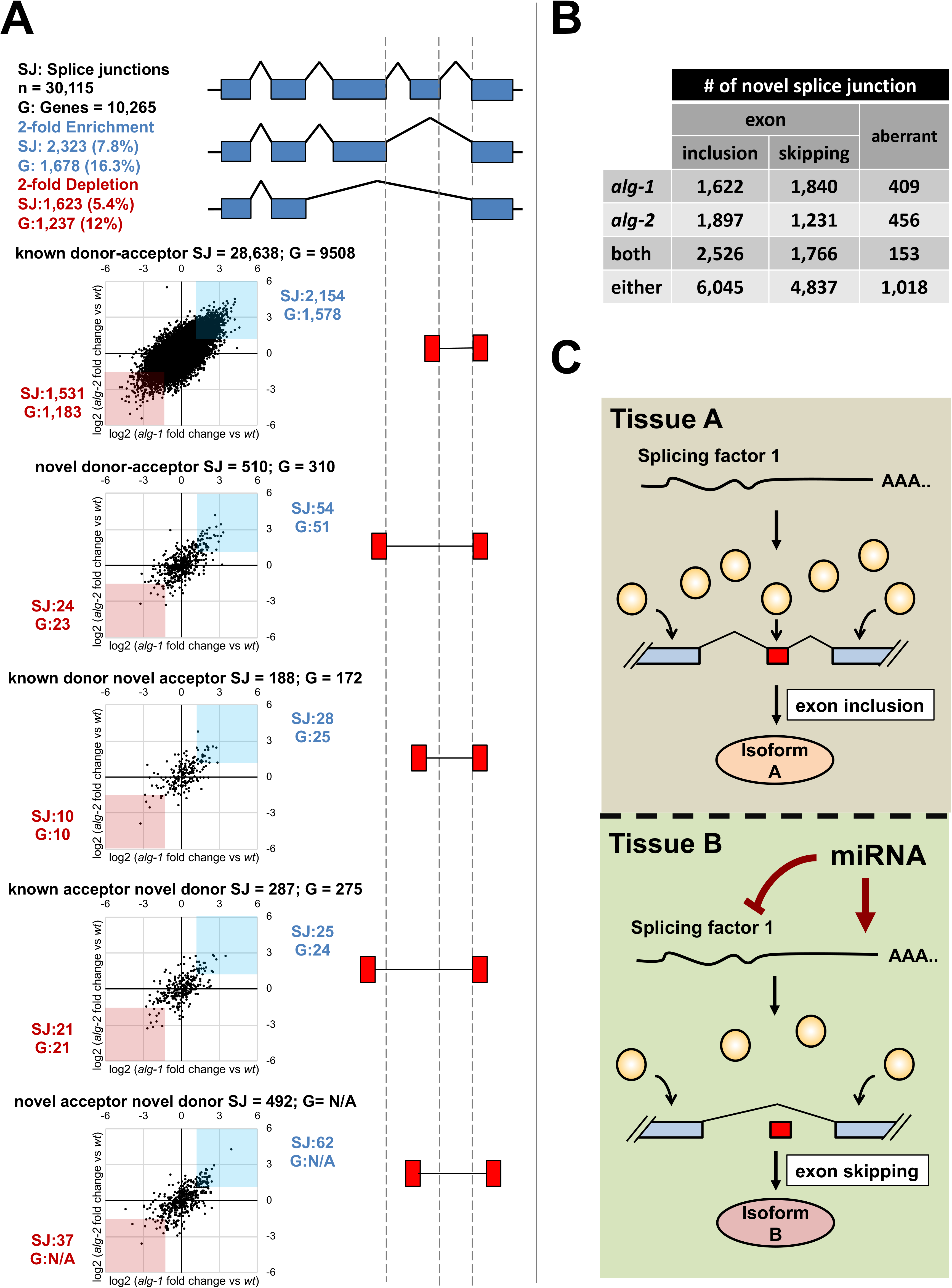
(A) Genome wide changes in splice junction usage in *C. elegans* strains deficient in the miRNA pathway. Analysis of splice junction (SJ) usage in miRNA deficient strains (*alg-1(gk214)* or *alg-2(ok304)*) re-annotated from Brown et al. 2017 (Brown *et al.* 2017). The graphs illustrate the changes in splice junction abundance for different types of splicing events. The x-axis represents the fold-change of the normalized number of reads for each splice junction, comparing the *alg-1(gk214)* strain to *wt*, while the y-axis represent the fold change obtained when comparing *alg-2(ok304)* to *wt*. Splice junctions with more than 2-fold enrichment in both strains are highlighted in blue, while splice junctions with 2-fold depletion in both strains are highlighted in red. The number of genes (G) with enriched or depleted splice junctions are indicated next to the graphs. There is a 2-fold change in ~13% of all the splicing events mapped in the knockout strains, effecting 3,301 genes, when compared to the N2 *wt* control. (B) The table summarizes the number of novel splicing events seen in *alg-1* and *alg-2* datasets. These are splicing events not observed in the N2 wt control, and indicate an increase in novel and aberrant splicing events in miRNA deficient strains. (C) A proposed role for miRNAs in the modulation of tissue-specific alternative splicing. The abundance of RNA splicing factors (yellow circles) dictates the splicing events in a given tissue A. The presence of a miRNA in Tissue B may lower the dosage of splicing factors resulting in tissue-specific alternative splicing.

## DISCUSSION

In this manuscript we have developed tools and techniques to identify tissue-specific miRNA targets and applied them to uniquely define the genes targeted by miRNAs in the *C. elegans* intestine and body muscle. We validated previous findings and mapped ~3,000 of novel tissue-specific interactions (Figure 2 and Table S1).

In order to perform these experiments, we have prepared worm strains expressing ALG-1 fused to GFP and expressed this cassette in the intestine and body muscle using tissue-specific promoters. We validated the ALG-1 expression (Figure S1), and the viability of our ALG-1 construct in *in vivo* studies (Figure S2). We have then performed ALG-1 immunoprecipitations in duplicate, separated the miRNA complex from their targets, and sequenced the resultant RNA using Illumina sequencing (Figure 1, Supplementary Figure S3). To confirm our results we validated few selected hits with expression localization studies in both tissues (Figure S5). Importantly, our ALG-1 pull-down results are in agreement with previous studies (Figure 2A, Figure S4, and Supplementary Figure S6), and are significantly enriched with predicted miRNA targets (Figure 2C).

The genes identified in this study overall match with the intestine transcriptome previously published by our group (81%) (Blazie *et al.* 2015; Blazie *et al.* 2017). Of note, only 56% of genes identified as miRNA targets in the body muscle match the body muscle transcriptome (Figure 2B). Perhaps, the remaining targets are genes strongly down-regulated by miRNAs in this tissue, leading to rapid deadenylation and mRNA degradation that make them undetectable using our PAB-1-based pull-down approach. Given the fact the body muscle transcriptome is significantly smaller than the intestine transcriptome, it may also be subjected to less regulation through miRNA. However, if we normalize the number of genes expressed in each tissue and study the proportion of the transcriptome targeted by miRNA, we still find significantly more regulation in the intestine (Figure 2A), suggesting that this tissue may indeed employ miRNA-based gene regulation to a greater extent.

We found a disparity in the proportion of each tissue-specific transcriptome targeted by miRNAs, with a notably larger proportion of genes targetted in the intestine. The majority of the targeted genes in our intestine pull-down IP are unique to the intestine and share only a handful of genes with our body muscle dataset (725 genes, 23% of the total intestine dataset) (Figure 2B). Conversely, very few genes are unique in the body muscle pull-down IP. The small pool of shared genes includes housekeeping genes that are most likely regulated similarly in both tissues. Of note, this minimal overlap between our tissue-specific datasets indicates that our ALG-1 pull-down is indeed tissue-specific with marginal cross-contamination.

Intriguingly, when we look at the miRNA population predicted to target the genes in our datasets as from MiRanda (Figure 2C Bottom Panels), we found an enrichment of known tissue-specific miRNA targets (Betel *et al.* 2010), which is in agreement with miRNA localization datasets (Alberti *et al.* 2018) (Figure S6). This in turn indicates that there is a tissue-specific miRNA targeting bias in *C. elegans*, with unique tissue-specific miRNAs targeting unique populations of genes.

Our experimental approach was designed for tissue-specific mRNA target identification, and unfortunately did not provide miRNA data. We assign tissue-specific targets to miRNAs relying primarily on prediction software and correlation to past-published datasets. These comparative approaches required conversion between genomic releases and data consolidation across different developmental stages and conditions, which may have add unwanted variability to our comparative analysis.

One of the most surprising findings of this study is that a large number of targets obtained with our tissue-specific ALG-1 pull-down are RBPs. 64% of the intestinal RBPs were found in our intestine ALG-1 pull-down, while 54% of the muscle RBPs were in our muscle ALG-1 pull-down. This result was unexpected given the small number of RNA binding proteins previously identified in the *C. elegans* genome (n = 887) (Tamburino *et al.* 2013), which amounted to only 4% of the total *C. elegans* protein coding genes. However, previous studies have hinted at a strong regulatory network between miRNAs and RBPs, as the 3′ UTRs of RBPs were found to contain on average more predicted miRNA binding sites than other gene classes (Tamburino *et al.* 2013).

Some of these RBPs in our top hits are well-characterized factors involved in the *C. elegans* fertilization and early embryogenesis but are not well documented in somatic tissues. For example, within our top 100 hits we obtained the genes *pos-1* and *mex-5* in the body muscle, and *gld-1* and *oma-2* in the intestine. We were surprised by these results, but at least in the case of *pos-1*, which is our top hit in our body muscle dataset, we validated its presence in the body muscle (Figure S5), suggesting a potential role for this and other RBPs outside the gonads.

RNA binding domain containing proteins are involved in many biological processes, and their role is not limited to RNA biogenesis (Tamburino *et al.* 2013). RBPs can bind single or double strand RNAs, and associate with proteins forming ribonucleoprotein complexes (RNPs). Longevity, fat metabolism, and development are all processes controlled by RNPs (Lee and Schedl; Masuda *et al.* 2009; Aryal *et al.* 2017), and in the context of miRNA regulation, the ability of miRNAs to control RBPs abundance and function allow for an increased control of fundamental cellular core processes. 234 RBPs uniquely detected as miRNA targets in the intestine, while 147 RBPs are shared between both datasets.

Within this intestinal dataset we mapped a surprising number of RBPs involved in RNA splicing (Figure 3B). We performed a literature search for known RNA splicing factors in *C. elegans*; out of the 72 total protein identified, 37 of them were detected at different level of strength in our intestine ALG-1 pull-down. In contrast, we do not observe this level of complexity in the body muscle, with only 9 RNA splicing factors identified in this dataset (Figure 3B).

*asd-2* and *smu-2* are well-known RNA splicing factors that induce exon retention in a dosage dependent manner (Spartz *et al.* 2004; Ohno *et al.* 2012), while *hrp-2* abundance lead to exon skipping (Kabat *et al.* 2009). Here we show that all three RNA splicing factors possess regulatory targets within their 3′ UTRs (Figure 4) that amount to ~40% silencing activity in the intestine (Figure 4). Although we do not know which miRNAs target the *asd-2* and *hrp-2* 3’UTRs, in Figure 5A and Figure S8 we show that the miRNA pathway influences splice junction usage by regulating these genes, and the depletion of miRNAs which target these RNA splicing factors by using sponge approaches led to defects in the alternative splicing pattern of downstream genes regulated by *asd-2* and *hrp-2*.

Interestingly, the miRNAs predicted to target most splicing factors were not found highly expressed in this study. *miR-85* and *miR-355*, the most abundant and tissue-restricted miRNAs identified, are only predicted to target less than 10% of all the RBPs found. This suggests that since miRNAs are highly reactive, the abundance of those involved in RNA alternative splicing may be tightly regulated in tissues, to make sure splicing events are properly executed.

Our genome-wide splice junction mapping effort in miRNA deficient strains shows similar trends of aberrant splicing of *unc-60*, *unc-52, lin-10* and *ret-1* (Figure 5B and Figure S9), and display an overall disruption of splicing events (~13.2% of all splice junctions mapped) (Figure 6A-B). Most of these defects are in known donor-acceptor splicing events, perhaps because RNA surveillance mechanisms may hide more severe disruptions.

Unfortunately, our *in vivo* approach does not reach the resolution needed to conclusively pinpoint the extent of the miRNA pathway in this process. In order to perform *in vivo* experiments, we used total RNA extracted from transgenic worms, and studied change in exon abundance occurring in a single tissue within a whole animal, which prevented us from reaching the same resolution obtainable with *in vitro* splicing experiments and mini-genes. In addition, the effects we observe are ameliorated by the presence of at least one functional Argonaute protein, which is able to compensate for the loss of the other. Knockout of the entire miRNA pathway is lethal in *C. elegans*, and while aberrant splicing may play a role in producing this phenotype, these activities are challenging to detect *in vivo*.

Taken together, our results support a role for miRNAs in regulating alternative splicing in the intestine, where their presence in a tissue-specific manner may lead to alteration of the dosage balance of RNA splicing factor, leading to tissue-specific alternative splicing (Figure 6C). MiRNAs are known to alter gene expression dosage, rather than induce complete loss of protein function (Wolter *et al.* 2017; Bartel 2018). On the other hand, many RNA splicing factors involved with constitutive and alternative splicing are ubiquitously expressed (Shin and Manley 2004), but are somehow able to induce tissue-specific alternative splicing in a dosage dependent manner. In this context, it is feasible that miRNAs may alter the dosage of RNA splicing factors, leading to tissue-specific alternative splicing (Figure 6C).

We have uploaded our intestine and body muscle miRNA target datasets into the 3’UTRome database (www.UTRome.org), which is the publicly available resource for the *C. elegans* community interested in 3’UTR biology (Mangone *et al.* 2008; Mangone *et al.* 2010). In order to provide a more comprehensive overview, we have also manually curated and included results from several available datasets including PicTar (Lall *et al.* 2006) and TargetScan (Lewis *et al.* 2005) miRNA target predictions, experimentally validated ALG-1 interaction (Zisoulis *et al.* 2010; Kudlow *et al.* 2012), tissue-specific gene expression and expanded 3’UTR isoform annotation data (Jan *et al.* 2011; Blazie *et al.* 2015; Blazie *et al.* 2017).

## Author Contribution

MM and KK designed the experiments. KK executed the experiments. ALS executed a portion of the experiments. HSS assisted with the experiments and the imaging of the *C. elegans* transgenic lines and performed the analysis in Figure S4. MM and KK performed the bioinformatics analysis and uploaded the results to the UTRome.org database. MM and KK led the analysis and interpretation of the data, assembled the Figures, and wrote the manuscript. All authors read and approved the final manuscript.

## Conflict of Interest

The authors declare that they have no competing interests.

### Acknowledgements

We thank Stephen Blazie for the cloning of the *alg-1* coding sequence used as the backbone for the development of the vectors used in this manuscript. We thank Dr. Honor Glenn and Dr. Jordan Yaron for their advice and assistance when using the confocal microscopes and image acquisition. We also thank Heather Hrach for insights and review of the manuscript. We thank Gabrielle Richardson, for maintaining the *C. elegans* strains used in this manuscript.

