## Supplementary Figures for "miRNA activity contributes to accurate RNA splicing in *C. elegans* intestine and body muscle tissues"

Additional File 1 for:

**This PDF includes:**

**Figures S1-S9**

**Table of Contents**

### Additional File 1

**Figure S1:** Validation of transgenic strains for miRNA-target identification…..…...3

**Figure S2**: Genotyping and brood size assay to validate *in vivo* the functionality of the GFP tagged ALG-1 construct...........................4

**Figure S3**: Summary of the sequencing results from our tissue specific ALG-1 immunoprecipitation..…………………………......5

**Figure S4:** Comparison of our tissue specific ALG-1 pull-down to previously identified *alg-1* RNA (Zisoulis et al., 2010)...................…6

**Figure S5:** Validation of selected hits detected in our tissue specific ALG-1 pull-down ……………………………………………………...7

**Figure S6:** Comparative analysis of our tissue specific ALG-1 pull-down to previously identified intestine and body muscle miRNA (mime-seq) (Alberti et al., 2018)..…………………………………………..……..8

**Figure S7:** Brood size assay to determine the efficacy of RNAi targeting splicing factors………………………………………………………………….9

**Figure S8:** *lin-10, unc-52 and ret-1* exon skipping events in the intestine (1)……10

**Figure S9:** *lin-10, unc-52 and ret-1* exon skipping events in the intestine (2).....11

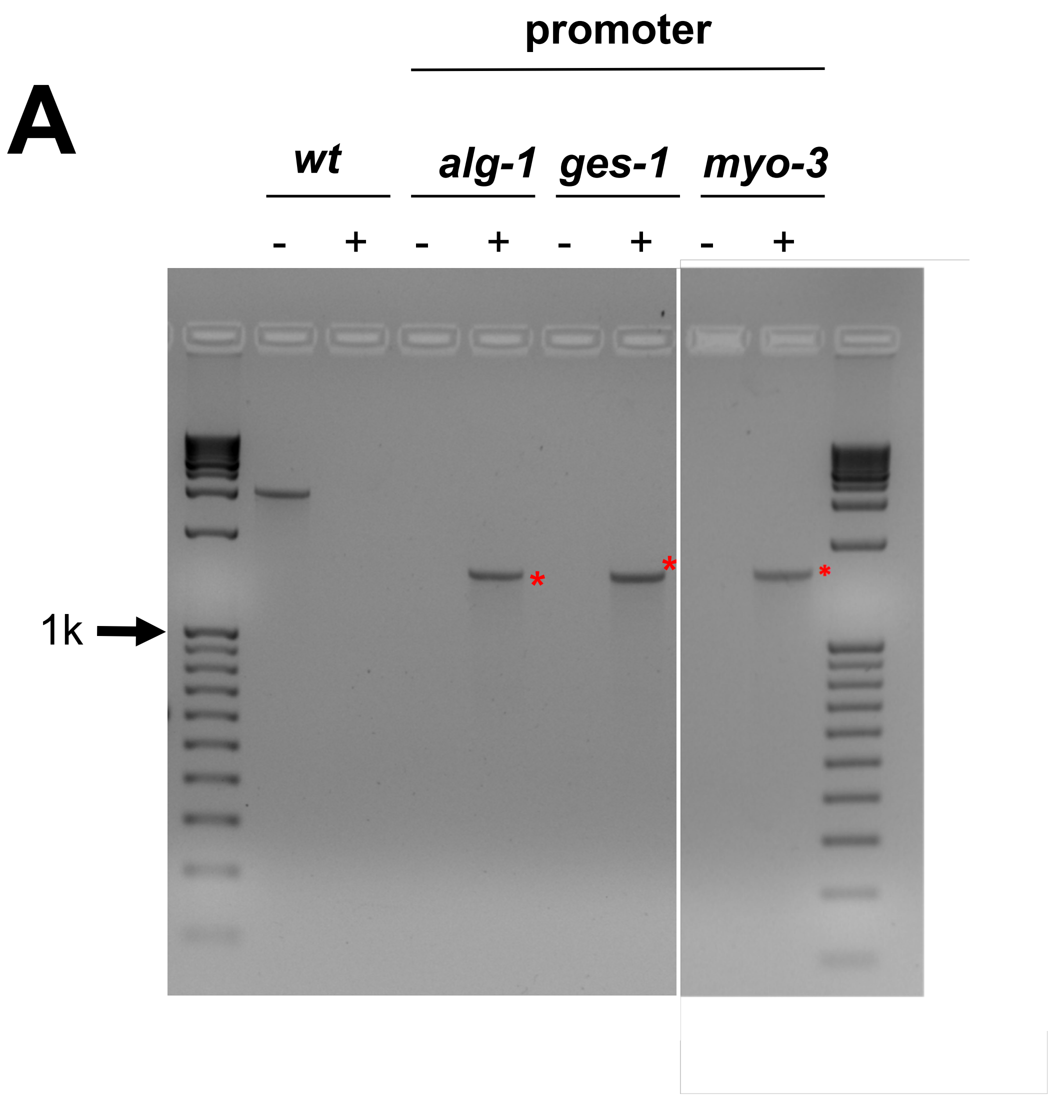


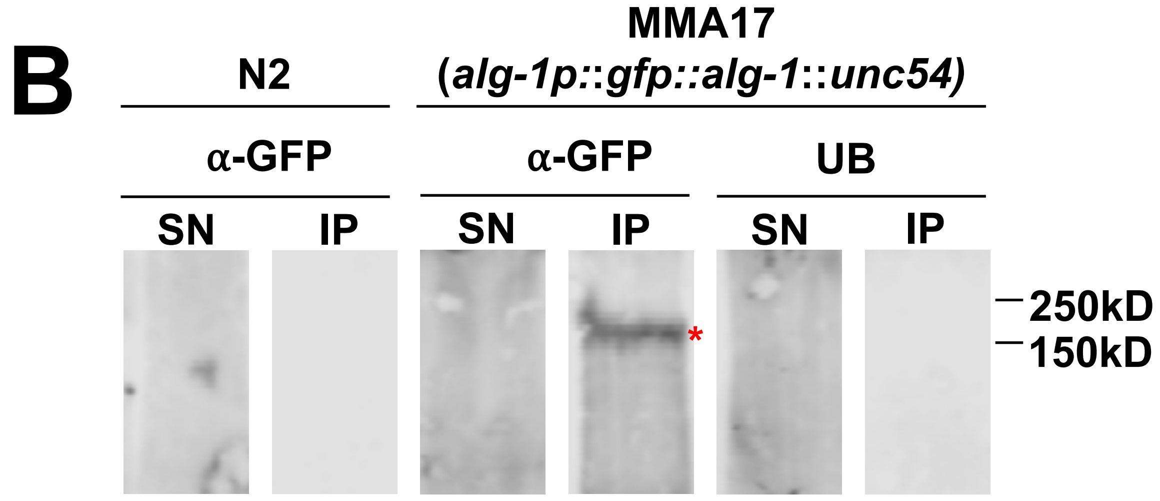


**Supplemental Figure S1:Validation of transgenic strains for miRNA-target identification.** A) Agarose gel electrophoresis of PCR products obtained from genomic DNA using single-worm PCR approach from transgenic *C. elegans* strains prepared in this study. Each strain was tested using primers that detect either the presence (+) or the absence (-) of the integrated MosSCI transgene. Red asterisks mark expected bands confirming integration of our constructs. B) Western blot experiments from *wt* N2 worms or endogenous ALG-1 pull-down construct performed using either α-GFP antibodies or unconjugated beads (UB) on immunoprecipitated protein obtained from each strain. (IP) immunoprecipitated protein, (SN) supernatant. Red asterisk marks the expected band.

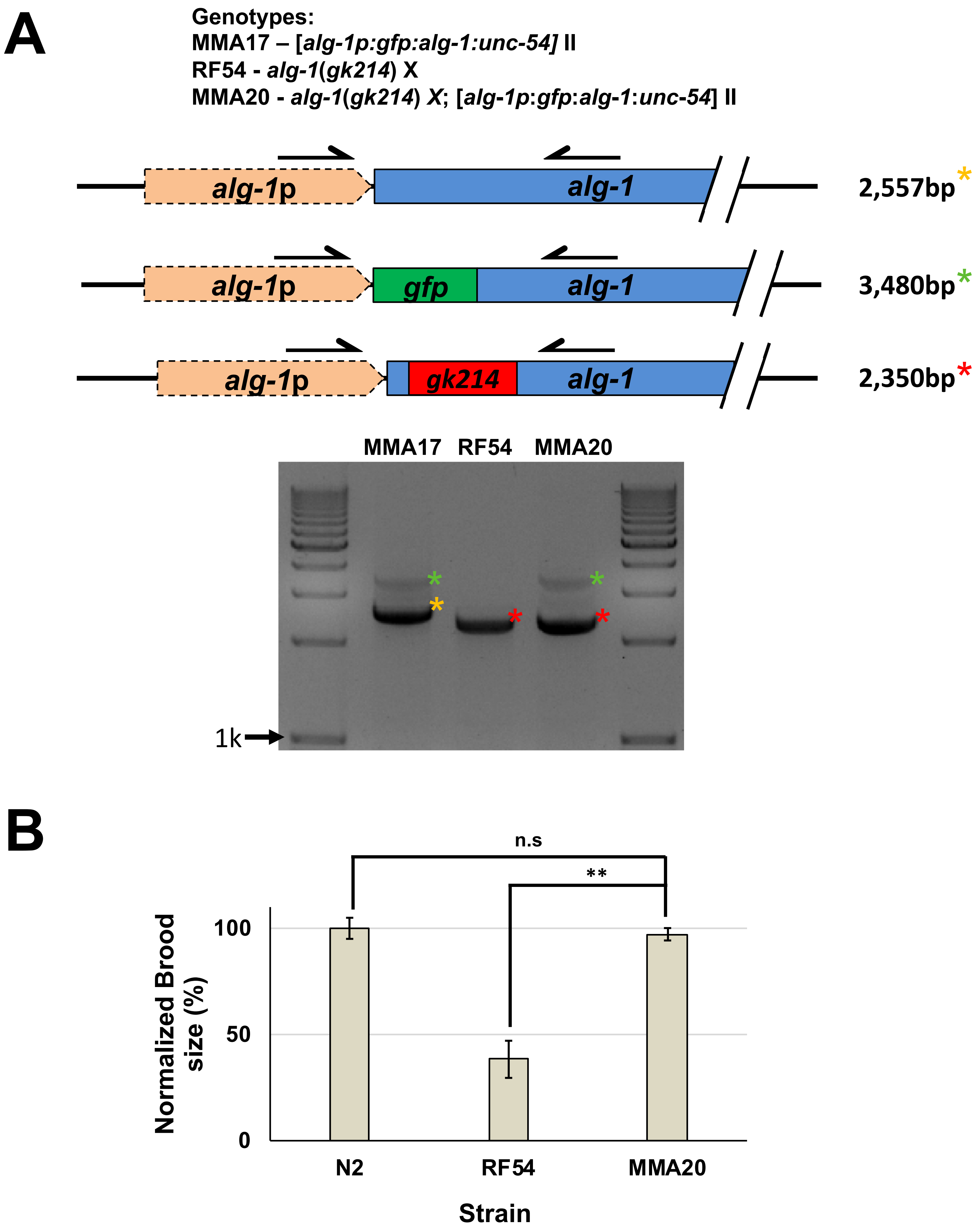


**Supplemental Figure S2: Genotyping and brood size assay to validate *in vivo* the functionality of the GFP tagged ALG-1 construct.** A) Agarose gel visualizing single worm genomic PCRs testing the presence of the *gk214* mutation before and after the cross. Yellow asterisks indicate the genomic locus with *wt alg-1* (2,557bp). Green asterisks indicate *gfp* tagged transgenic *alg-1*. Red asterisks indicate the presence of the *gk214* mutation (2,350bp). In the MMA20 strain we re-introduced our cloned *alg-1* to the RF54 strain, which lacks of endogenous *alg-1.*
B) Results from the brood size assay testing fertility. The loss of fertility seen in the *alg-1* knockout strain RF54 is rescued in the MMA20 strain which expresses our GFP tagged transgenic ALG-1, suggesting that our ALG-1 pull-down construct is functional and mimic endogenous *alg-1* activity.
Student T-test. ** p<0.01. Error bars indicate the standard error of the mean.

**A**

| Sample | | Sequencing reads Total (#) | Mapped (#) | Mapped (%) |
|---|---|---|---|---|
| ***alg-1p*** | | 23,277,135 | 18,292,992 | 78.6 |
| **Intestine** | Experiment (*rep 1*) | 31,424,437 | 23,241,554 | 73.9 |
| | Replicate 2 (*rep 2*) | 23,522,056 | 17,861,605 | 75.9 |
| **Body muscle** | Experiment (*rep 1*) | 24,092,664 | 21,327,095 | 88.5 |
| | Replicate 2 (*rep 2*) | 25,347,564 | 22,062,749 | 87.0 |
| N2 | | 21,894,026 | 19,378,018 | 88.5 |

**B**

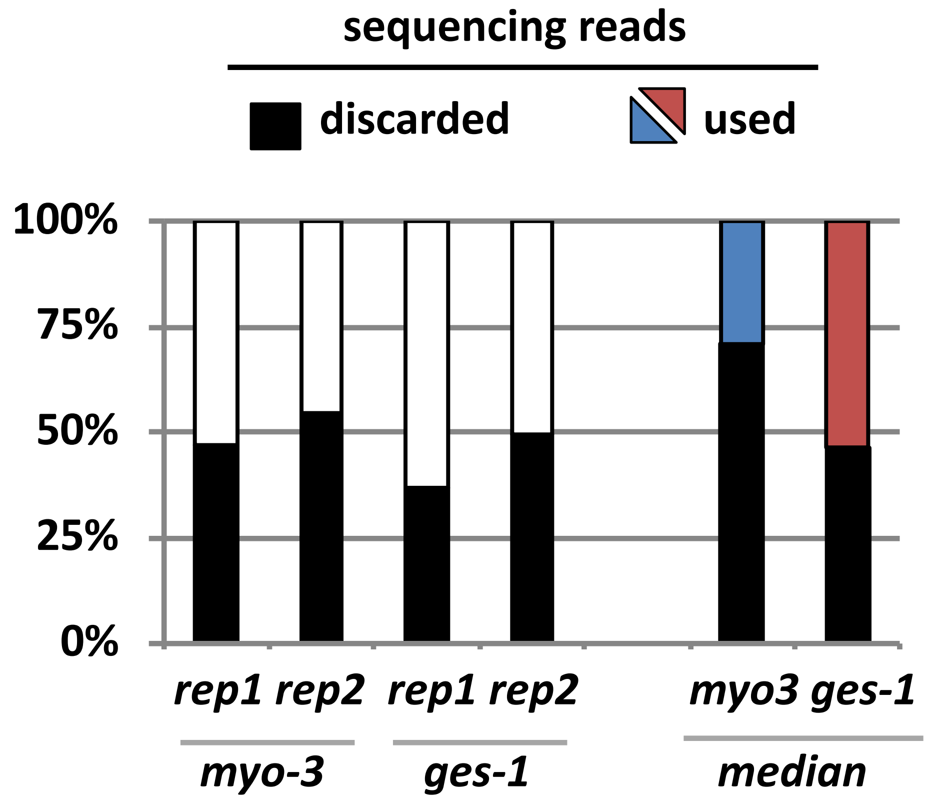


**C**

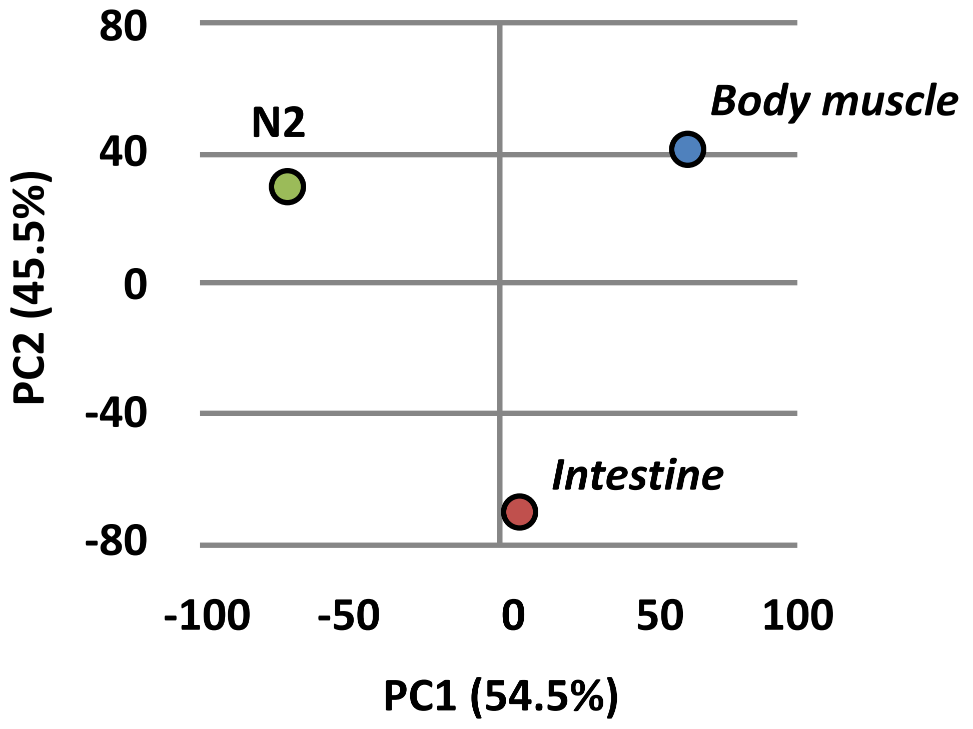


**Supplemental Figure S3: Summary of the sequencing results from our tissue specific ALG-1 immunoprecipitation.** A) Overview of the sequencing reads obtained for each immunoprecipitation. Each sample was sequenced in duplicate with biological replicates. We obtained at least 15M reads each dataset mapped to the *C. elegans* genome (WS250). B) In this study we have used only the top ~50-75% positive hits produced by the Cufflinks algorithm. . Blue; Body muscle. Red; Intestine C) Principal Component Analysis (PCA) between the median *fpkm* values among replicates shows differences between tissues within our datasets. Each dot represents a median experiment and replicate.

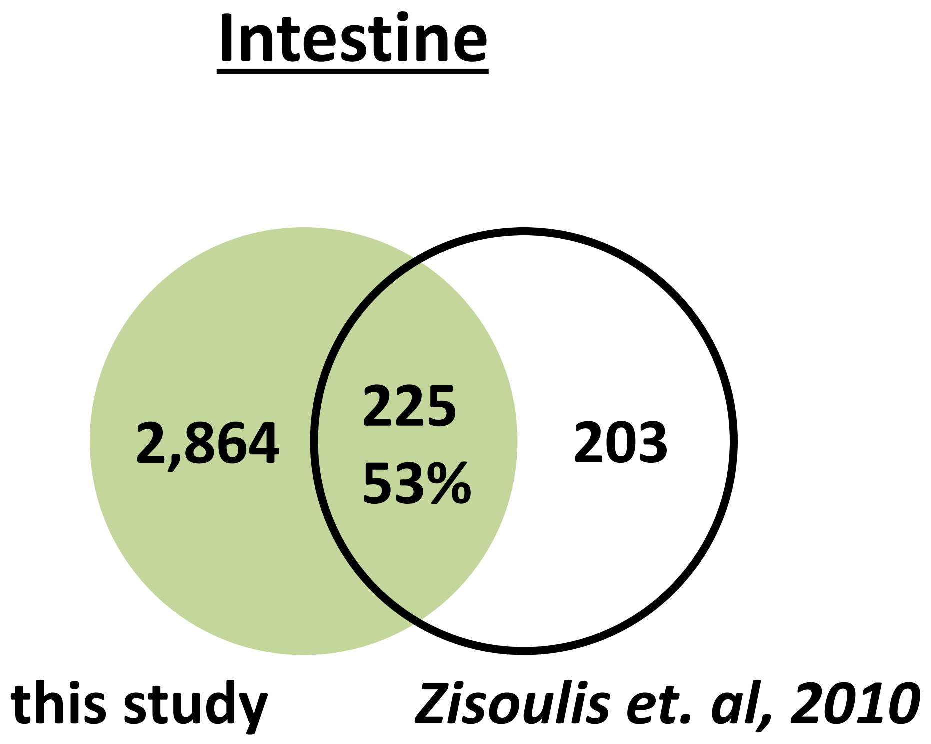


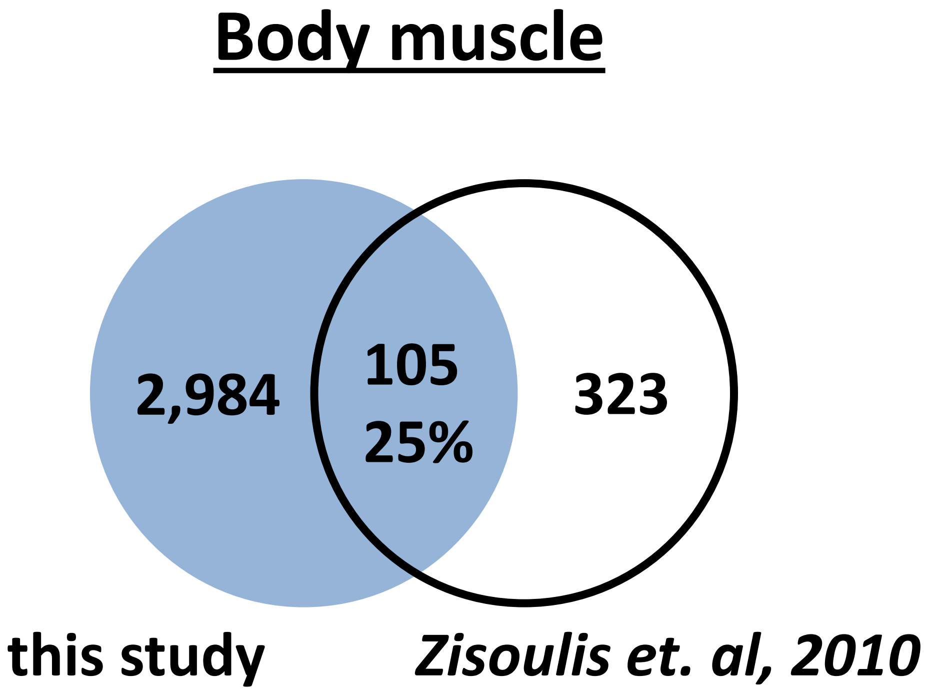


**Supplemental Figure S4**: **Comparison of our tissue specific ALG-1 pull-down to previously identified *alg-1* RNA (Zisoulis et al., 2010).** Venn diagram showing the comparison of hits between our datasets and a comprehensive RNA pull down of miRNA targets.

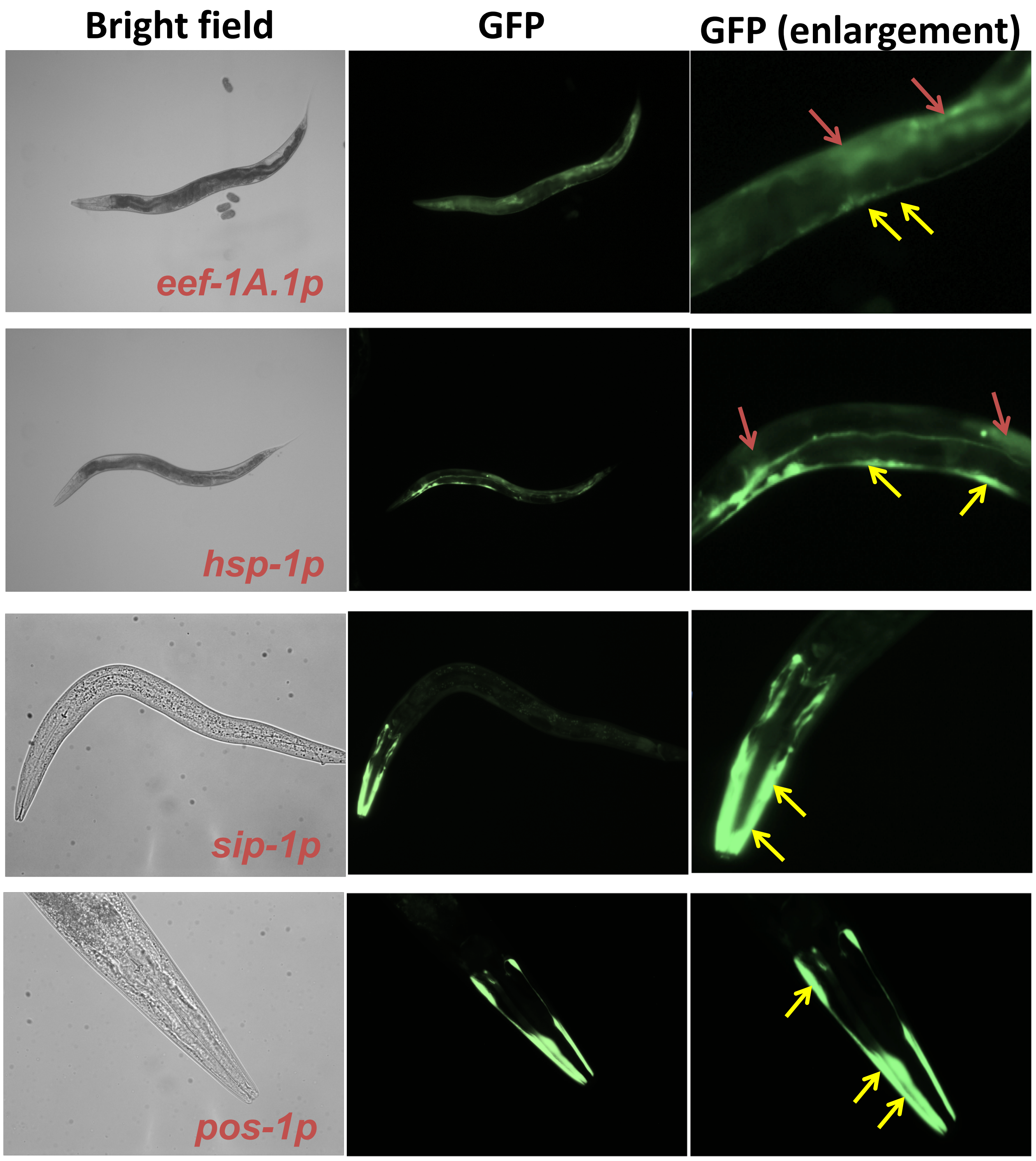


| Name | Intestine | | Body muscle | |
|---|---|---|---|---|
| | GFP Detected | Validated | GFP Detected | Validated |
| *sip-1* | ** | yes | *** | yes |
| *eef-1A.1* | *** | yes | *** | yes |
| *pos-1* | - | no | *** | yes |
| *hsp-1* | *** | yes | *** | yes |

**Supplemental Figure S5: Validation of tissue localization selected hits detected in our tissue specific ALG-1 pull-down.** We have cloned selected promoters from genes identified in our tissue specific ALG-1 pull-down, and fused it to GFP reporters to validate the expression localization pattern. Except for one case, we detected strong GFP expression in the expected tissues. Red arrow mark intestine expression, Yellow arrow mark body muscle expression. Putative expression index: - not detected; * low expression; ** expressed; *** strong expression

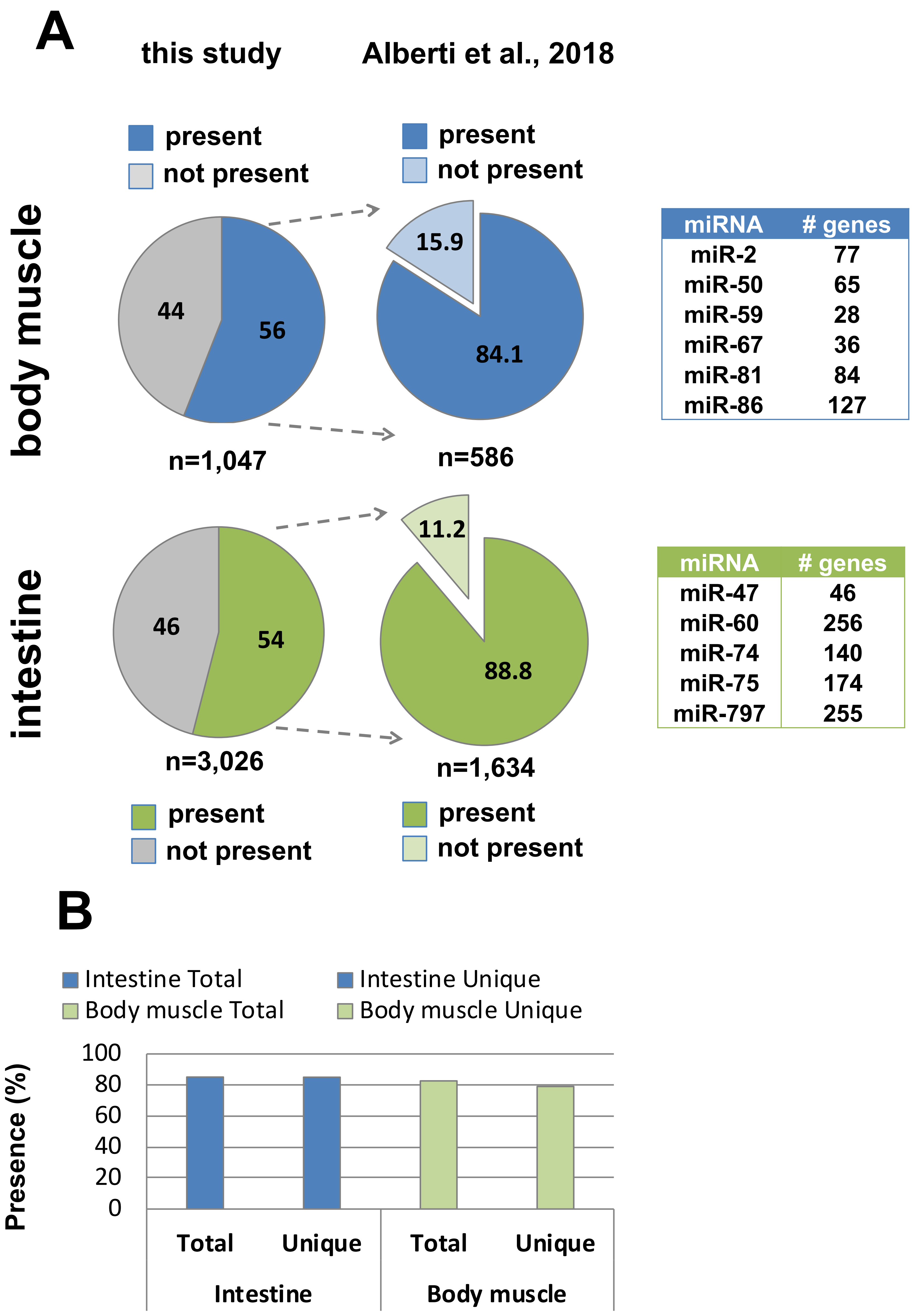


| miRNA | # genes |
|---|---|
| miR-2 | 77 |
| miR-50 | 65 |
| miR-59 | 28 |
| miR-67 | 36 |
| miR-81 | 84 |
| miR-86 | 127 |

| miRNA | # genes |
|---|---|
| miR-47 | 46 |
| miR-60 | 256 |
| miR-74 | 140 |
| miR-75 | 174 |
| miR-797 | 255 |

**Supplemental Figure S6**: **Comparative analysis of our tissue specific ALG-1 pull-down to previously identified intestine and body muscle miRNA (mime-seq) (Alberti et al., 2018).** A) The left pie chart shows the percentage of genes identified in our study with predicted miRNA binding sites as for MiRanda algorithm. The right pie chart shows how many of those genes contain a seed region for intestine or body muscle miRNAs identified in Alberti et al., 2018. The table shows how many genes identified in our study are targeted by unique tissue-specific miRNAs identified in Alberti et al., 2018. B) Percentage of total and tissue-restricted (unique) intestine or body muscle ALG-1 targeted genes identified in our study with seed regions for miRNAs identified in Alberti et al., 2018

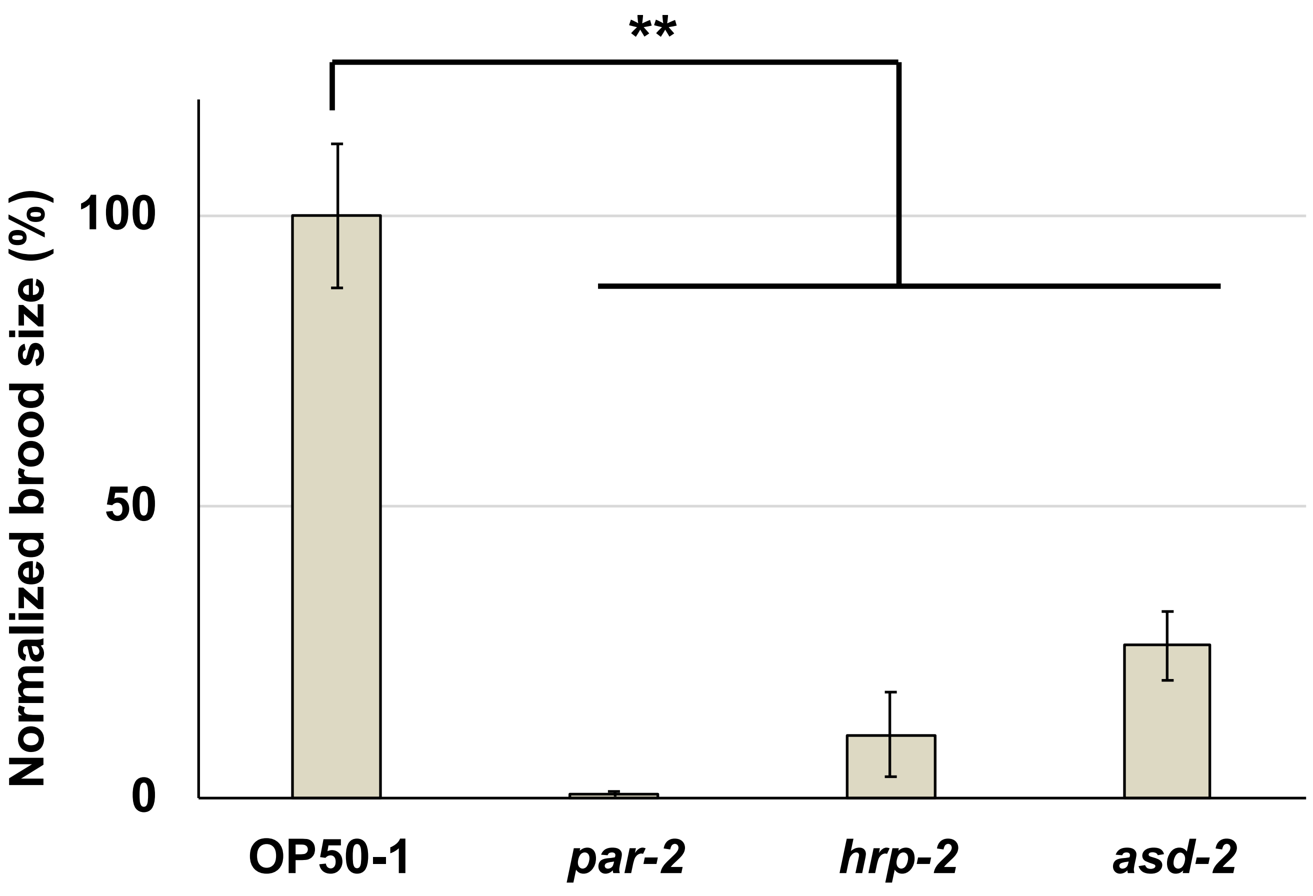


**Supplemental Figure S7: Brood size assay to determine the efficiency of RNAi targeting *hrp-2* and *asd-2*.** RNAi performed against the gene *par-2* (positive control) produced almost no hatched larvae as expected. A decrease in brood size was observed when RNAi was performed for the splicing factors *hrp-2* and *asd-2* indicating the knockdown of these genes. **p<0.01 n=5

#### Supplemental Figure S8

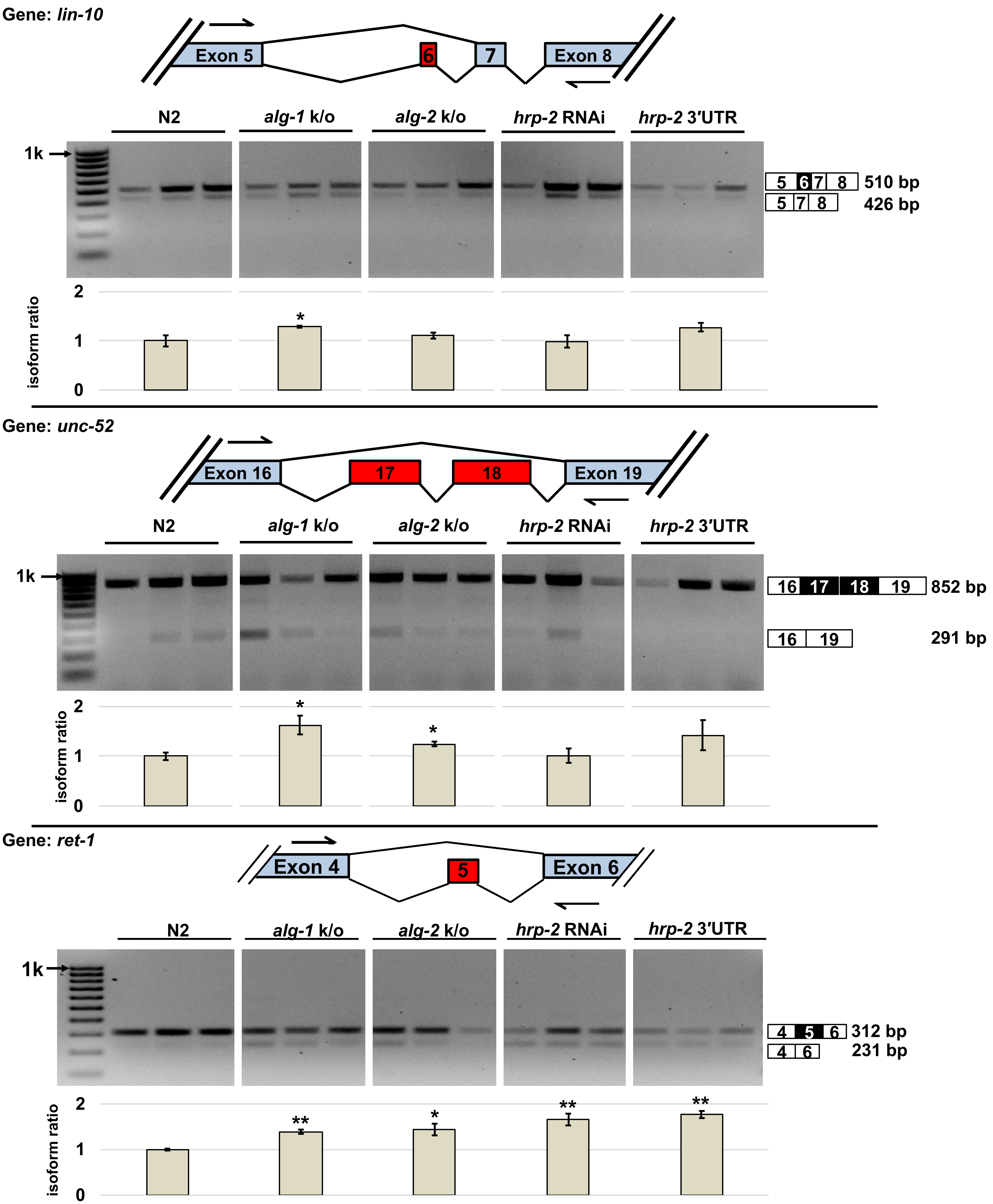


**Supplemental Figure S8: *lin-10, unc-52 and ret-1* exon skipping events in the intestine (1).** Top panels: The *lin-10, unc-52* and *ret-1* gene models are shown to illustrate the alternatively spliced exons (red) and the primer pairs used in the RT-PCR experiments to detect intestine specific alternative splicing modulated by miRNAs. Bottom Panels: Agarose gels visualizing RT-PCRs performed in triplicate on independent RNA extractions to test changes of alternative splicing of individual genes. *p < 0.05; **p <0 .01

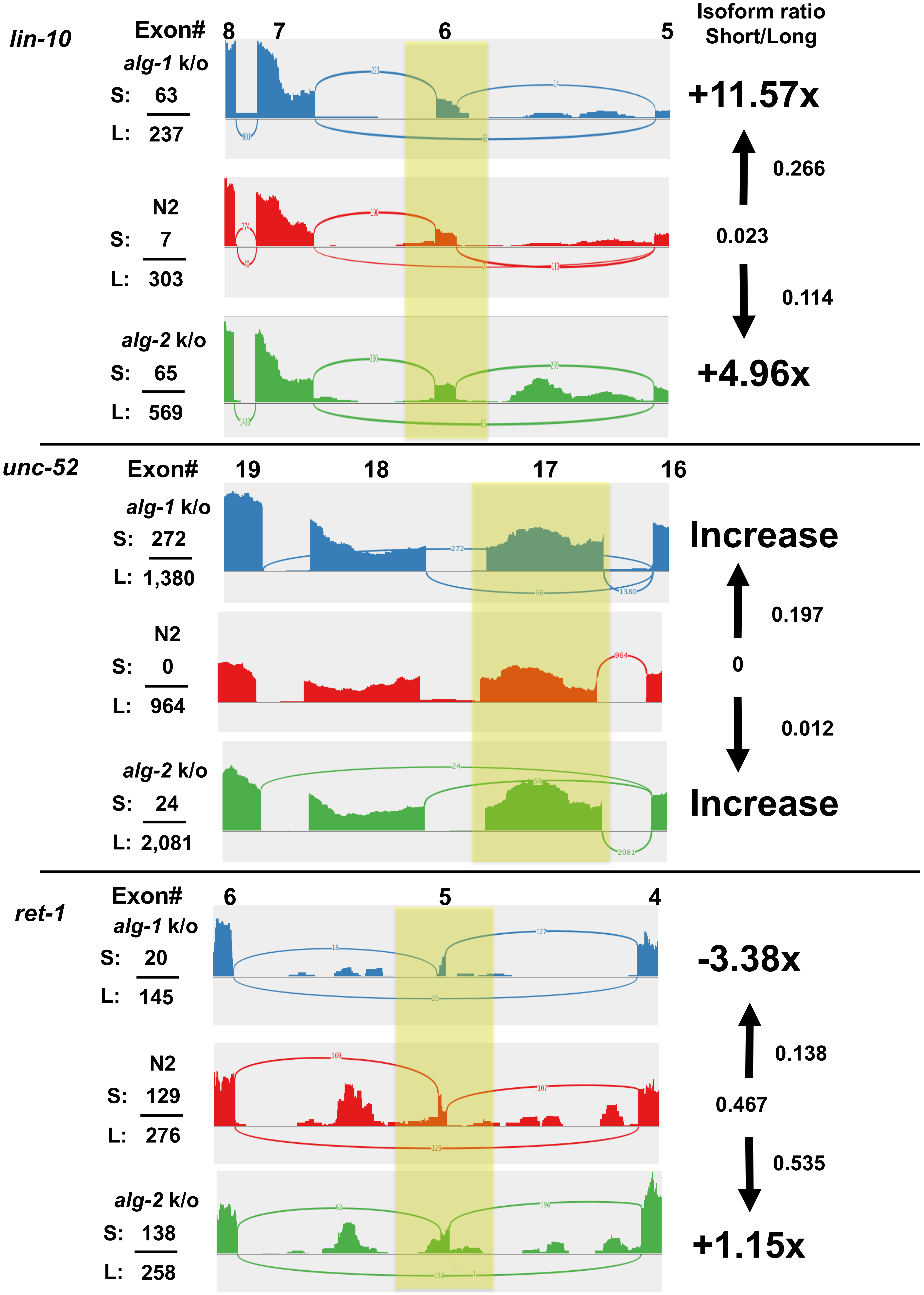


**Supplemental Figure S9: *lin-10, unc-52 and ret-1* exon skipping events in the intestine (2)** Sashimi plots generated by re-analyzing RNA-seq data from Brown et.al 2017. Replicates for each strain were combined and analyzed with the TopHat bioinformatic package. The plots illustrate the number of reads mapped for each splice junction. Top and middle panels: The splicing events for *lin-10* and *unc-52* showed an increase of exon skipping occurring in both miRNA deficient strains. Lower panel: *ret-1* exon 5 skipping increases slightly in *alg-2* but shows increased exon inclusion in *alg-1* knockout worms. Yellow boxes highlight alternatively spliced exons analyzed in this figure S: total reads for the short isoform; L: total reads for the long isoform.
